# Thinking outside of the box: Refining rat housing to improve welfare

**DOI:** 10.64898/2026.04.29.721812

**Authors:** Carly I. O’Malley, Emilie A. Paterson, Halimatou Tambadou, Erik Moreau, Olufemi Ekundayo, Jukka Puoliväli, Chereen Collymore, Patricia V. Turner

**Affiliations:** Global Animal Welfare & Training, Charles River, Wilmington, MA, USA; Discovery Services, Charles River, Kuopio, Finland; Laboratory Animal Medicine and Surgery, Charles River, Senneville, QC, Canada; Department of Pathobiology, University of Guelph, Guelph, ON, Canada

**Author notes:** Corresponding author (CIO).

**Keywords:** 3Rs, nonaversive handling, housing, behavior management, rats, animal welfare

## Abstract

Standard rat housing may impede species-typical behaviors and impact rat welfare and research outcomes. This research investigated the effects of housing on behavioral and physiological outcomes of rats through the use of modified large animal cages for housing, and was conducted in two studies. Study A: 70 Sprague Dawley (SD) rats (34 males, 36 females; 5 wk old) were randomly assigned to standard polycarbonate shoebox cages (C: 733.9cm^2^) or modified stainless steel primate cages (T: 10,416cm^2^) for 18 days. In Study B: 48 SD rats (24 males, 24 females; 7.5 wk old) were held in T housing for 90 days to assess long term impacts. All rats received gentle handling for 15s 3x/week. Rats were assessed for body weight, anxiety-like behavior in an elevated plus maze, response during a voluntary human approach test, and overall home cage behavior, posture, and space usage. Data were analyzed using generalized linear mixed models, with sex and treatment as fixed effects, and cage as the random effect. The results of study A suggest that the modified large animal cages (T) had positive impacts on rat behavior and welfare. T rats were less anxious (*P*=0.038) and more active (*P*<0.0001) and explorative (P=0.0003) compared to C rats. In both groups, activity levels declined towards the end of the 18-day study period (*P*<0.0001). For study B, similar patterns were observed, with rats becoming more inactive (*P*<0.0001) over 90 days. However, rats spent significant time on elevated shelves in T housing, which increased throughout the study (P<0.0001), suggesting continued use of the resources the housing provided. In both studies, there were no differences in latency to approach humans (P>0.05), but T rats spent less time in contact with human handlers, suggesting differences in motivation to interact with humans that should be explored further.

## Introduction

The 3Rs is a well-accepted framework that emphasizes continuous improvement of research animal welfare. Refinement, one of the 3Rs, is described as minimizing pain and distress of research animals [1], advancing animal welfare and gaining a better understanding of the role animal welfare plays in scientific outcomes [2]. Within research environments, animals play a key role in the development of needed health interventions for human and veterinary medicine, such that ensuring positive animal welfare is necessary for research reproducibility and translatability [3]. Rats are a commonly used biomedical species and despite domestication, rats are highly motivated to perform species-specific behaviors such as burrowing, climbing, and stretching vertically [4]; however standard rat housing in shoebox cages restricts their ability to perform these behaviors [5].

Improving housing environments for research rats is not only beneficial for rat welfare, but also for reducing overall number of animals used and improve the quality of the science. A recent systematic review by Cait et al. (2022) highlights the importance of an appropriate housing environment on the health of research rodents. Standard housing for rodents limits their ability to perform natural behaviors and can affect their physical and mental health [6]. For mice and rats of both sexes, conventional housing resulted in increased susceptibility and morbidity related to cancer, cardiovascular disease, stroke, depression, and anxiety [6]. Mortality rates were also increased. The findings of this study emphasize the importance of appropriate and thoughtful housing on the validity, translatability, and ethics of housing rodents in conventional cages as standard practice [6] as well the need to carefully describe housing environments in published research using rodents for context [7]. Previous work on rat housing has suggested that rats prefer a larger housing enclosure with more resources [8] and that it is beneficial behaviorally and physiologically.

Research animal welfare is an important social and scientific topic and animal behavioral management programs should aim to reduce negative emotional states in animals as well as finding ways to promote positive experiences and emotional states. Housing considerations should include for animal and human health and safety and behavioral interactions [9], as well as the behavioral diversity that animals exhibit based on the resources provided [10]. The ability to display natural behaviors and postures is an important outcome of the housing environment and this is being reflected increasingly in regional guidance, such as the rat guidelines by the Canadian Council of Animal Care (2020), which indicates that housing must provide enough height to allow rats to stand in a vertical posture (up to 30 cm for fully grown male rats), enough space to allow rats to be socially housed and to walk, jump, run, and play, and sufficient resources to allow rats to avoid light and open space, build nests, create microenvironments, forage, gnaw, and climb [11]. The CCAC guidelines recommend approximately 800 cm^2^ of floor space per rat depending on size of the rats, suggesting 1500 cm^2^ of floor space for four 100 g weanlings to allow them sufficient space for rough-and-tumble play, and 1500 cm^2^ for two 600 g rats [11]. At the time of this study, there were few commercially available housing options for rats that meet these guidelines.

Besides housing and resources, welfare considerations also need to include human-animal interactions. Anecdotally, it is often thought that rats housed in larger, more complex environments will be more difficult to work with. Rats have an innate fear of humans [5,12] but they respond well to gentle handling and habituation for husbandry and study procedures [13]. Regular gentle handling has positive effects on rat behavior and stress levels, including decreased anxiety [13-15], depression [15], and reduced fear towards humans [16,17], improved learning and memory [18], and more reliable results in behavior tests [16] compared with rats without regular gentle handling. When human-interactions are positive, rats will actively seek out human contact by pressing a lever in an operant test chamber to gain access to a human [19]. However, despite evidence of the benefits of regular gentle handling for rats by habituating them to contact with people for husbandry and scientific procedures, few institutions have a standard protocol in place incorporating this practice. If rats are not habituated to human presence or the interactions are not positive, rats show a stress response even to simple tasks such as cages being removed from racks and lids opened [20]. Therefore, it is in the best interest of the animals and the research outcomes to implement regular gentle handling prior to study start to habituate rats to interactions with humans, especially in more complex housing systems.

The objective of this research was to investigate the impacts of housing enclosure type on the behavior and physiology of research rats. This research was conducted across two studies. The first study (Study A) aimed to compare behavioral and physiological parameters between rats housed in different housing environments, including standard housing and primate cages that had been modified to meet the behavioral needs of rats, as well as investigating the impact of regular gentle handling on response to humans based on housing environment. It was hypothesized that rats in the resource-enhanced housing would a) display less anxiety-like behaviors, due to previous work investigating the effects of improved in-cage resources on rat behavior and welfare [6], b) show more vertical postures due to this being a species-typical behavior and there being more freedom of movement in this housing [4,5], and c) there would be no difference in response to humans due to implementation of regular gentle handling. The second study (Study B) aimed to evaluate the long-term effects of housing rats in modified primate cages by assessing changes in behavior over time as rats habituated to the housing. Based on the initial results of Study A, it was hypothesized that a) rat activity levels would decrease over time in the modified primate changes as rats habituates to the housing, b) that these rats would weigh less than rats in standard housing due to increased activity in the additional space, and c) that rats’ response to humans wouldn’t be affected by housing due to regular habituation.

## Materials and Methods

All rat experimental protocols were approved by the Charles River Senneville Institutional Animal Care and Use Committee (#30816). The facility maintains a certificate of Good Animal Practice provided by the Canadian Council on Animal Care (CCAC) and is accredited by AAALAC, International.

### Study A

#### Animals

A total of 70 Crl:CD(SD) Sprague Dawley rats (34 males and 36 females) were obtained from Charles River (Kingston, NY) and housed at a Charles River pre-clinical safety assessment facility in Senneville, QC, Canada in June and July 2021. Rats were approximately 5 weeks old at the start of the study (median weight: 111.7 g) and were bred and raised at the site. The study length was 18 days. At study start, the rats were evaluated for health and received individual identification using Stoelting nontoxic animal markers (Wood Dale, IL, USA). Animals were weighed the day prior to transfer onto study and again at days 7 and 14.

Rats received *ad libitum* access to pelleted Lab Diet Certified CR Rodent Diet 5CR4 (St. Louis, MO, USA) and had *ad libitum* access to municipal tap water treated by reverse osmosis and ultraviolet irradiation from water bottles. The animal housing rooms were maintained between 21-25°C, 40-70% relative humidity, and with a 12:12 light:dark cycle. Prior to study start, rats were housed in in pairs or trios with littermates. The housing was as described below for control rats. The rats received minimal handling prior to study start, including daily checks and body weights.

#### Housing

There were 2 housing treatments in Study A. Rats were randomly assigned to housing treatment in same sex groups balanced by body weight. To do so, animals were listed by animal ID, sex, and weight. The animals’ weight was cut at the median to group animals of a similar weight together. Animals within the same weight group were then assigned to treatment by entering the treatment type into a list randomizer (random.org/lists). Then treatment groups were allocated to their cages by entering the cage IDs into the list randomizer. If littermates were allocated to the same cage, they were reassigned to ensure all animals were unfamiliar. Cage change and/or cleaning occurred on days 7 and 14. The two housing treatments were standard housing for research rats in polycarbonate cages (control, C) and large animal cages modified for rats that had more space allowance and resources (treatment, T).

##### Control (C)

Polycarbonate cages (Lab Products LLC, Aberdeen, MD, USA) were used (48.3 cm L x 35.6 cm W x 20.3 cm H, for a total floor space of 1,719.5 cm^2^) and contained 170 g of hardwood chip bedding (Envigo 7090 Teklad Sani-Chips®, Indianapolis, IN, USA), 1 tunnel (Bio-Serv Rats Retreats^TM^, Certified, Flemington, NJ, USA), one half of a Nylabone© (Neptune City, NJ, USA) per rat, a rotating in-cage resource (1^st^ week: Envigo Teklad 7979C.CS Certified/Irradiated Diamond Twists, Indianapolis, IN, USA; 2^nd^ week: timothy hay, Bio-Serv Certified, Flemington, NJ, USA), and one 4 g nest puck (Bed-r’Nest puck, The Andersons, Delphi, IN, USA) per rat (Fig 1). Animals were socially housed with 3 rats per cage across 8 cages (24 rats total in C housing).

**Fig 1.**
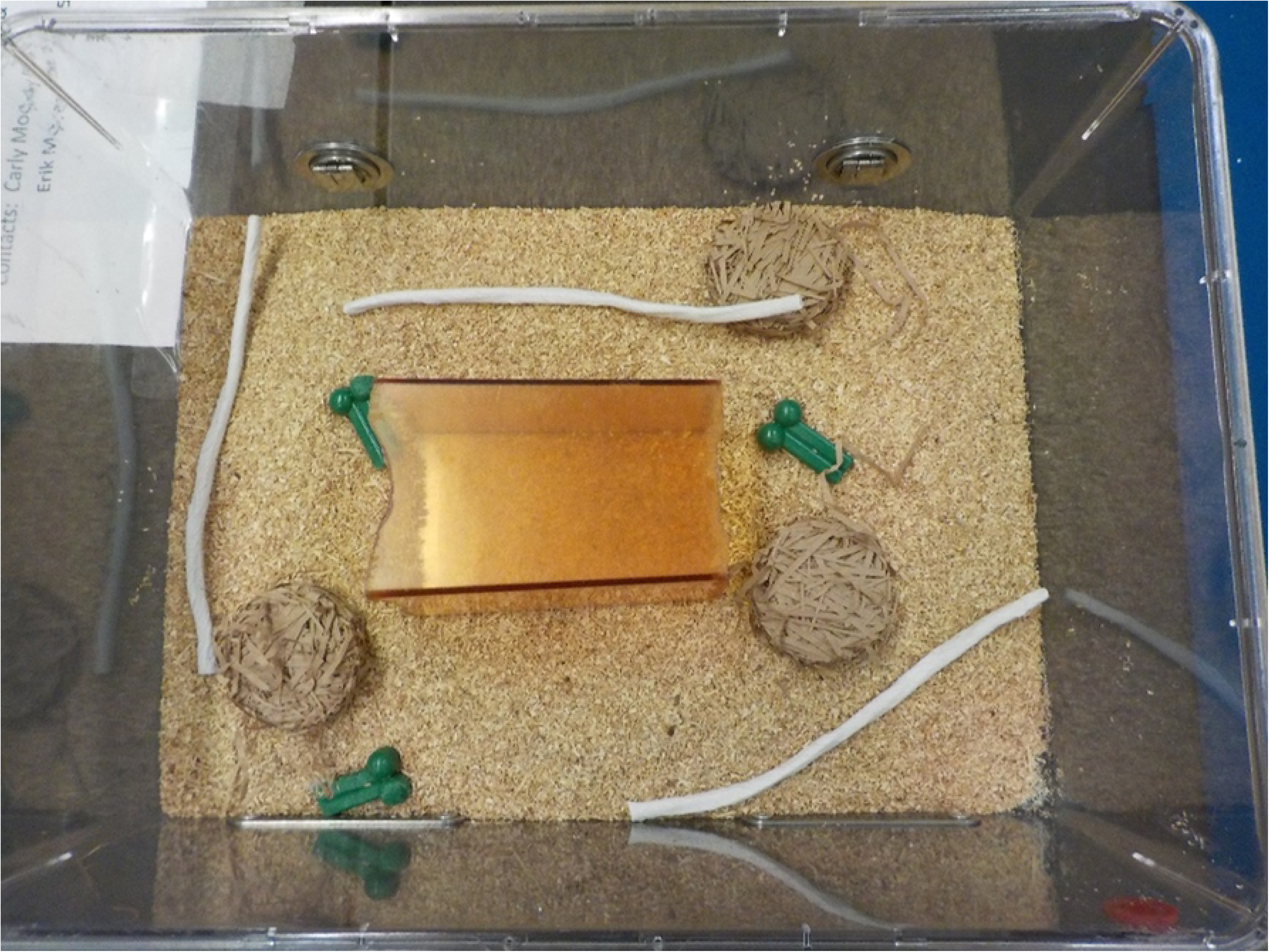
Standard rat housing (48.3 cm L x 35.6 cm W x 20.3 cm H; 1,719.5 cm^2^ of total floor space) with 170 g beta chip bedding, three half Nylabone©, three Diamond Twists, and three 4g nest pucks. Animals were socially housed with three rats per cage.

##### Modified cages (T)

Stainless steel primate cages (original manufacturer unable to be identified; see Fig 2a) (81 cm L x 83 W x 93 cm H) were adapted to house rats. In-house modifications included replacing the solid side and back walls with 1.3 cm stainless steel wire mesh with square holes. The bottom floor grate was removed and sealed to permit rats direct access to the removable floor pan. The cage door was replaced with clear Lexan high density polyethylene (HDPE) allow for video recording. White and clear HDPE was used to build two interior tunnels spanning the depth of the cage with dimensions 61 cm L x 24 cm W x 16.5 cm H, with holes on the side and top to permit rats multiple entry and exit points. A clear panel was placed on the inner sides of the shelves to allow for better animal viewing for the video recordings (see Fig 2). A hammock was added and made of stainless-steel wire mesh with dimensions of 51 cm L x 15 cm W. The hammock was suspended between the two side walls of the cage. The total floor space encompassed 10,416 cm^2^. within the cage, rats were provided with 1530 g of hardwood chip bedding (Envigo 7090 Teklad Sani-Chips®, Indianapolis, IN, USA), 1 tunnel (Bio-Serv Rats Retreats^TM^, Certified, Flemington, NJ, USA), 1 half Nylabone© (Neptune City, NJ, USA) per animal, a rotating resource item (1^st^ week: Envigo Teklad 7979C.CS Certified/Irradiated Diamond Twists, Indianapolis, IN, USA; 2^nd^ week: timothy hay, Bio-Serv Certified, Flemington, NJ, USA), two 4 g nest pucks per rat (Bed-r’Nest puck, The Andersons, Delphi, IN, USA), and one shredded tissue placed on the hammock (Fig 3). Rats were socially housed with 5-6 rats per cage across 8 cages (46 rats total in T housing).

**Fig 2.**
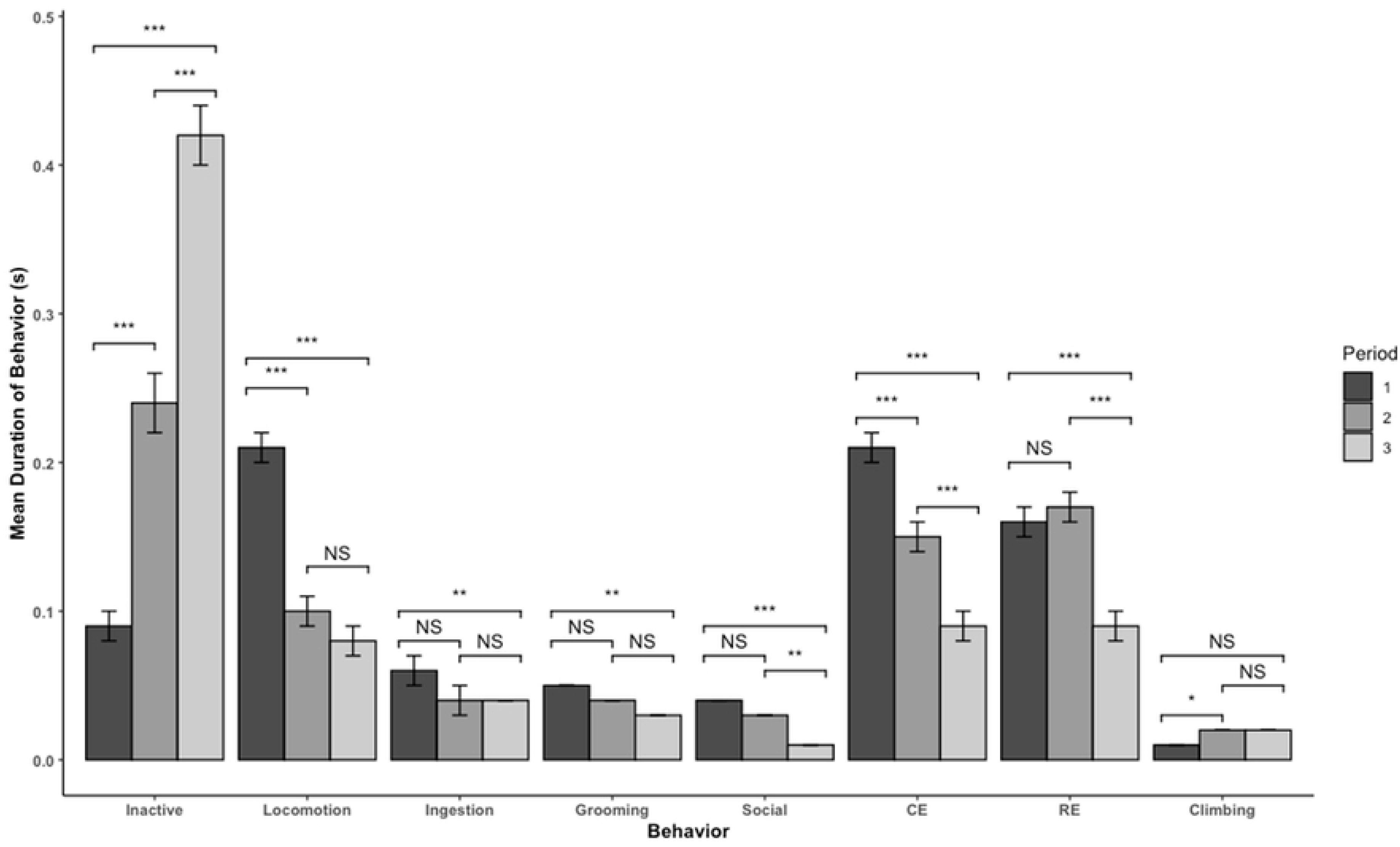
a.) original primate cages. b.) modified cages for rats (91.4 cm L x 91.4 cm W x 77.5 cm) pictured on the top cage. Cages were modified by replacing the side and back walls with 1.3 cm wire mesh with square holes (Fig 2c), sealing the bottom of the cage to allow access to the removable pan, replacing the doors with clear HDPE, placing two HDPE shelves (61 cm L x 24 cm W x 16.5 cm H), and placing a hammock (stainless steel wire mesh; 51 cm L x 15 cm W). The total floor place of the modified cages was 10,416 cm^2^.

**Fig 3.**
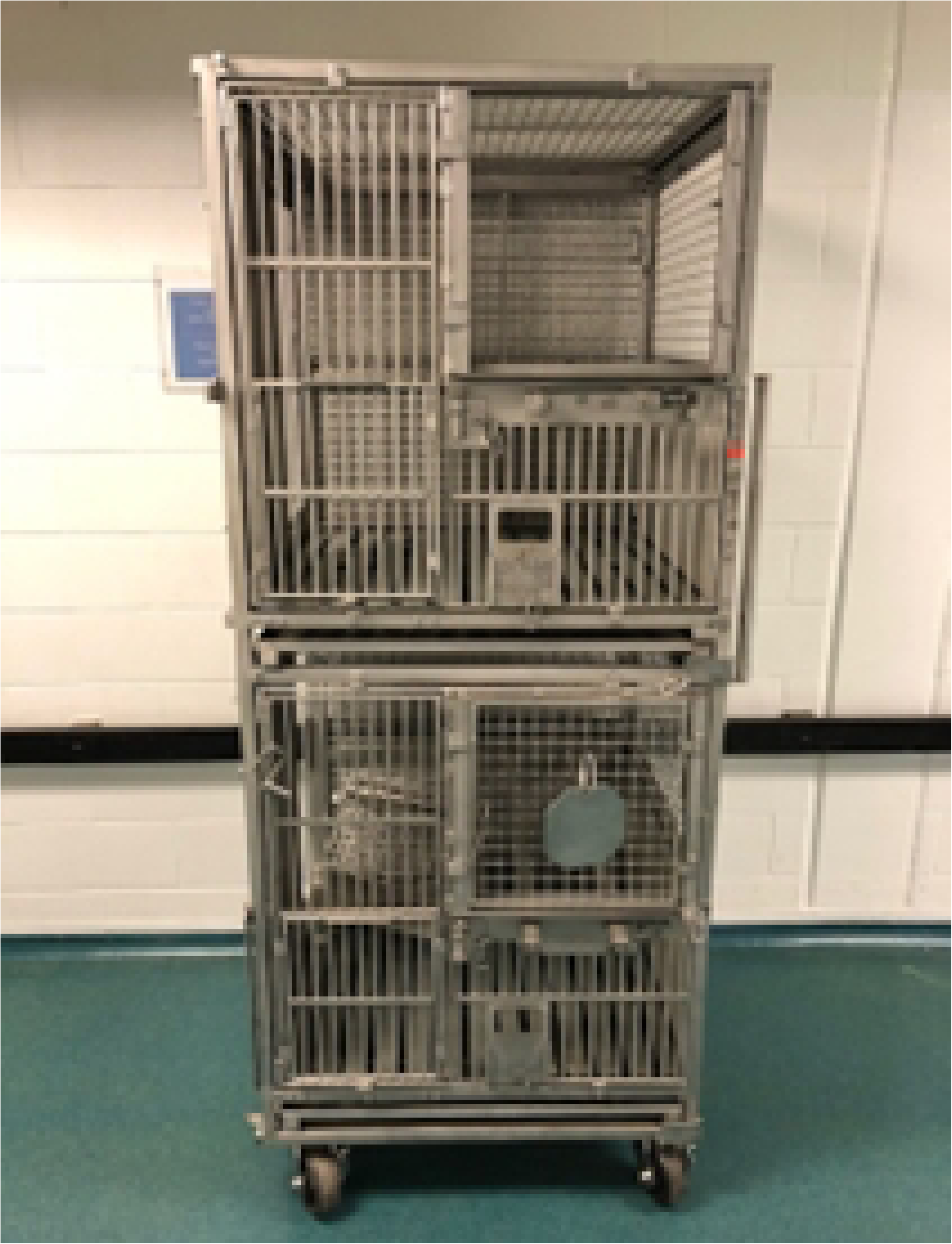
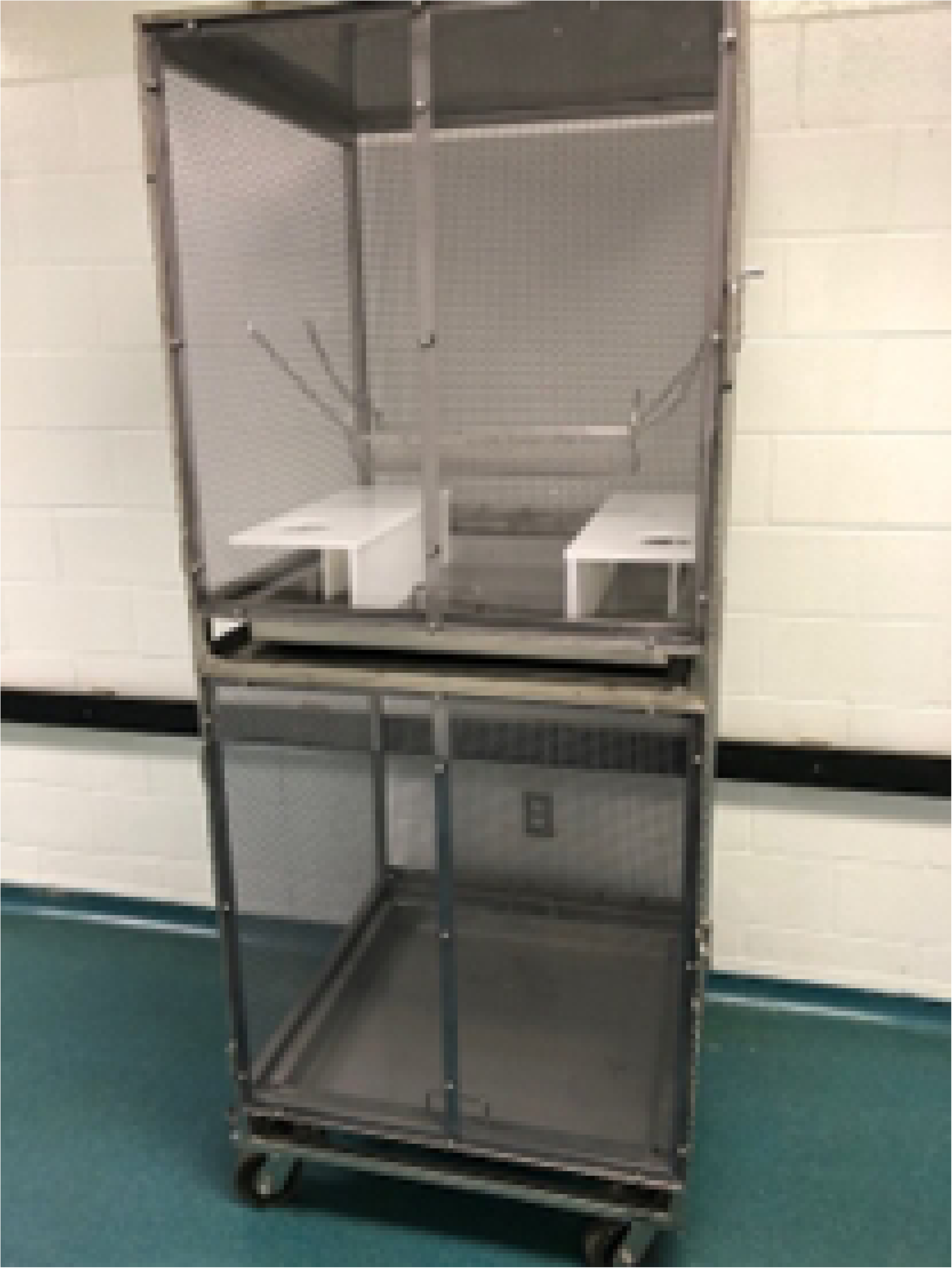
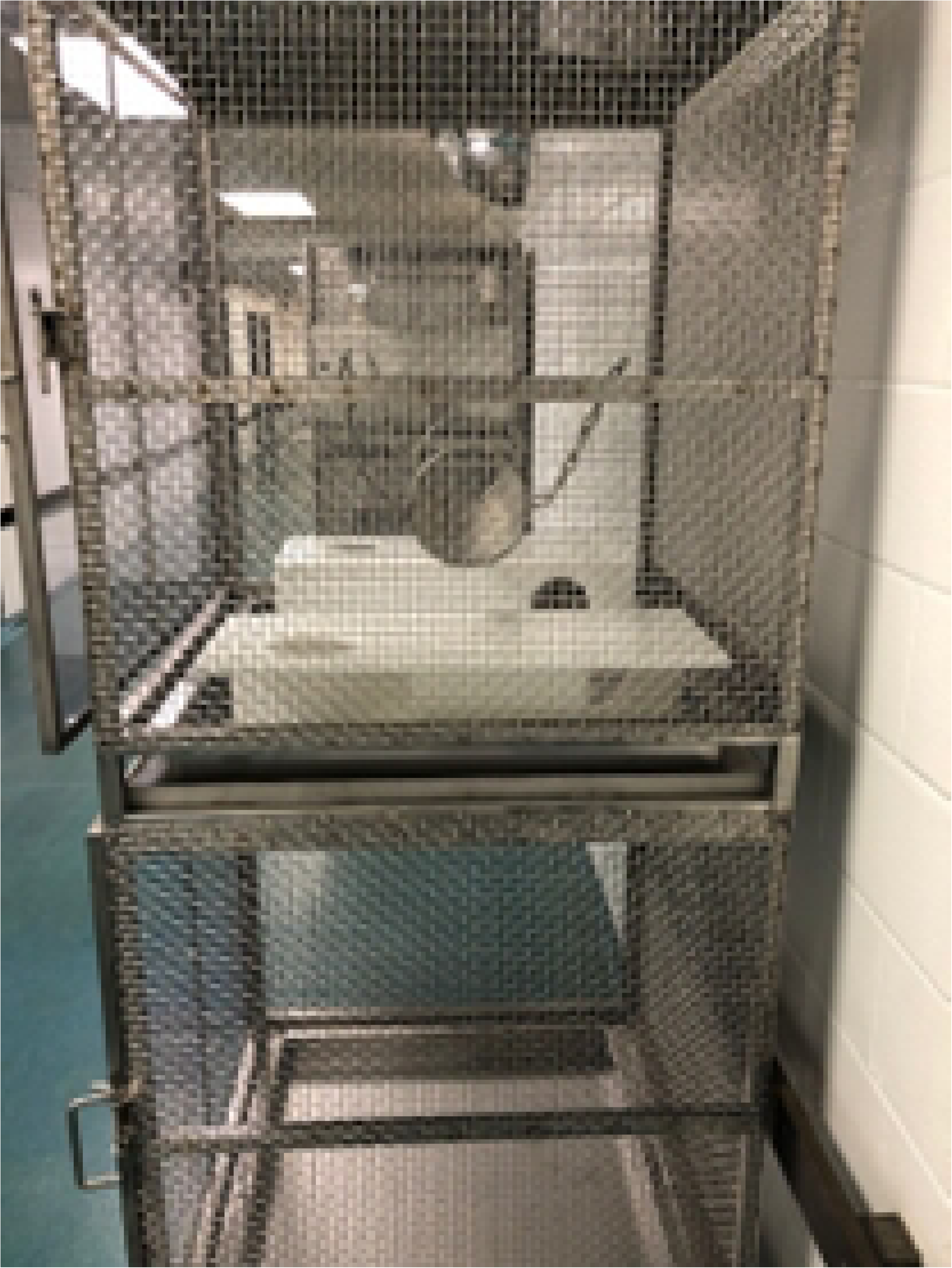
The final version of the modified cages, which included 2 HDPE shelves (61 cm L x 24 cm W x 16.5 cm H), a wire-mesh hammock (51 cm L x 15 cm W), 1530 g of beta chip bedding, 1 tunnel, one Nylabone© half per animal, one Diamond Twist per animal, two 4g nest pucks per animal, and one shredded tissue. Rats were housed 5-6 per cage.

#### Gentle Handling

Non-aversive body handling was always used (scooping from under the rat with one hand) to remove animals from cages and enclosures. All rats received regular gentle handling at a rate of 15 s per session approximately every other day for the first 13 days of the study (a total of 7 handling sessions). There were 2 handlers for this study (1 male and 1 female). Each handler interacted with 1 sex of animals for each handling session. The sex was selected at random for the first handling session then alternated throughout the rest of the sessions to maintain balance across animal sex and handler gender to account for possible differences in animal response towards male and female handlers [21] and differences in attitudes towards animals between genders [22]. For the gentle handling sessions, a fleece mat (30 cm L x 20 cm W; VetBed Canada, Smithers, BC,) was clipped onto the front of the handlers’ lab coat. For housing treatment C, a cage of rats was placed on a table, opened, and rats greeted. For housing treatment T, rats were transported from the cage using a transport box (spare standard rat cage: 48.3 cm L x 35.6 cm W x 20.3 cm H) on a cart. The transport box contained corncob bedding (Envigo 7092-7097 Teklad corncob bedding, Indianapolis, IN, USA). The transport box bedding was not changed between cages, but separate boxes were used for each sex. The transport box was placed on a table. One rat was scooped up at a time, then the handler sat, and set a timer for 15 s. Rats were handled gently and placed close to the mat clipped onto the lab coat. As part of the gentle handling, the rats were stroked along the head, body, tail, and limbs. After 15 s, rats received a treat (Cheerios©, General Mills, Minneapolis, MN, USA) and were placed back into the home cage (C) or into a separate transport box (T). All rats underwent the same procedure until all the rats in the same cage/enclosure had been handled, after which T rats were returned to their original enclosure. After 4 handling sessions with the above procedure, the gentle handling sessions included mock restraint for blood collection from the tail vein. Cages were randomized for handling but here was no fixed order for handling the rats in each cage or enclosure.

#### Behavior Tests

To test the effects of housing on anxiety in rats, an elevated plus maze (EPM) was used. The EPM consisted of 2 open arms (no sides/walls) and two closed arms (with sides/walls) made of black polycarbonate (Lexan) ¼ in thick. Anxiety was measured by examining the number of entries into the open vs closed arms. More entries or time spent in the open arms is associated with reduced anxiety whereas more entries or time spent in the closed arms is associated with increased anxiety [23,24]. The dimensions of the EPM were 110 cm total L, 45 cm L each arm x 10 cm W of each arm and the enclosed arms were 40 cm H. The EPM test occurred on day 15. Rats were tested individually and placed into the center of the apparatus facing the open arm opposite of the handler (the handler always stood at the same location). Rats were video recorded in the maze for 5 min. The order of testing was selected at random at the cage level.

To test the effects of regular gentle handling on human-rat relationships, a human approach test (HAT). For housing treatment C, a cage of rats was placed on a table. For housing treatment T, rats were placed into the transport box, which was then placed on a table. A novel person was used for the HAT but the same person did both tests and this handler also performed the blood collection on day 17. The person placed their left hand into the cage palm down so the hand was touching the bottom of the cage floor and the hand remained there for 60 s. The HAT was conducted twice, immediately before blood collection and immediately after on day 17. The order of testing was randomized at the cage level.

#### General Condition

Body weight (g) was collected on the day prior to the study start, and on days 7 and 14. Rats were weighed individually. Rats were also observed daily for signs of general condition throughout the study.

#### Blood Collection

Blood collection occurred on day 17. Both handlers from the study restrained the rats for blood collection. After the first HAT, individual rats were picked up by each handler (one rat per handler) and the handler sat in a chair with the VetBed on their lap. The handler restrained the rat gently for blood collection while a second person (the novel person from the HAT) collected blood from the lateral tail vein. A 25 g needle was used to prick the lateral tail vein to collect a drop of blood. A drop of blood was allowed to fall directly on a glucometer test strip (Bayer Contour Next EZ, Leverkusen, Germany). If blood was not forthcoming after 2 needle pricks, the rat was returned to the cage. Blood glucose levels were recorded for each rat in mg/dL. The order of the blood collection was the same as for the HAT, which was selected at random at the cage level.

#### Behavior Scoring

Rat behavior was continuously recorded in home cages throughout the study period using Reolink (RCL-410-5mp; Wilmington, New Castle, Delaware, USA) cameras. The elevated plus maze was also recorded using the same cameras. The human approach tests and blood collection were recorded using camcorders (JVC GZ-E200BU; Walnut, CA, USA). Behavior was scored using Noldus Observer XT 15 (version 15.0.1200; Leesburg, VA, USA). For all behavior scoring, there was a single observer per behavior measure (i.e., only one observer scored HAT, only one observer scored home cage behavior, one observer analyzed the EPM videos) for a total of 3 observers. Intraobserver reliability was assessed by comparing 10 videos scored twice for each measure of behavior. Greater than >80% reliability was deemed acceptable; however all intraobserver reliability reached >90% agreement. Order of videos watched was randomized by entering the list of video file names into a list randomizer (random.org/lists) and the observer was blinded to cage number and sex but could not be blinded to housing treatment for home cage recordings. For EPM, HAT, and behavior during blood collection, the observer was blinded to housing treatment, cage, animal number, and animal sex.

A 24-hour time budget was determined using instantaneous scan sampling, scoring the number of rats engaged in each behavior at the start of every hour for the 18-day study period. Behavioral time budgets were summarized for each behavior per day and for hour of the day. These data were also used to determine active periods for more in-depth observation where rats were observed for a 10 min duration at 5:00am every day, where duration of time (seconds) spent in each behavior, posture, and location was scored. The ethogram for behaviors scored is provided in Table 1. Postures included standing vertically, standing horizontally, lying down, and sitting. Location for the C cages included front of cage, back of cage, or tunnel. Location for T cages included: under left shelf (LH), under right shelf (RH), in tunnel (TB), on the floor (but not under the shelves; FL), on top of left shelf (OLH), on top of right shelf (ORH), on the hammock (HA), or on the walls (WA). In addition to active social behaviors scored from the ethogram, social proximity, defined as time spent in the same location as cage mates, was determined using instantaneous scan sampling by counting the number of rats in each location (corrected by total number of rats in the cage) at the top of every hour for the 18-day study period. For rats in T housing, it was confirmed if rats were within a body length of each other if in the same location, mainly to account for the location of walls, where rats may be climbing on the walls but not physically near one another. Confirmation of social proximity was done by a single observer.

**Table 1:** Ethogram used for scoring rat behavior in home cage. Behaviors were scored as mutually exclusive.

| Behavior | Definition |
| --- | --- |
| Inactive | Rat is not moving. Rat can have eyes open or closed. |
| Locomotion | Rat is actively moving around the cage, including walking, running, or climbing. |
| Ingestion | Rat appears to be eating or drinking (i.e., head is in the food bin or near the water sipper). |
| Grooming | Rat is licking or scratching itself. |
| Social behavior | Rat is engaged in non-aggressive interactions with conspecifics such as grooming or sniffing. |
| Aggression | Rat is engaged in behaviors such as wrestling, pinning, biting, or chasing with conspecifics, either as the giver or receiver. |
| Cage-directed exploration | Rat is interacting with cage fixtures or cage walls, such as sniffing or chewing. |
| Resource-directed exploration | Rat is interacting with the resources provided in the cage (i.e., tunnels, Nylabones, nest pucks, bedding) by sniffing, chewing, digging, or pawing. |
| Abnormal behavior | Rat is bar biting or self-chewing. |

To determine how rat behavior changed throughout the study period, data was split into 2 periods: period 1 (days 1-9), period 2 (days 10-17). On day 2 of video recording, camera malfunction resulted in no video recordings at 5:00 pm, therefore no data from day 2 was included in the analyses, resulting in 8 days of observation for period 1 and 8 days of observation for period 2.

An elevated plus maze (EPM) was used as a measure of anxiety-like behaviors and general locomotory activity. For the EPM, rats were scored using video tracking and movement analysis software (Noldus EthoVision XT, Leesburg, VA, USA). Rats were scored on the number of entries into the closed and open arms and the duration of time spent in the closed and open arms. Anxiety-like behavior was calculated as duration of time spent in open arms (s)/total time spent in open and closed arms (s). General locomotory activity was calculated as total number of arm entries into open and closed arms [23-26].

For the HAT, rats were scored on latency to touch the hand (s) and duration of time spent in contact with the hand (s).

During blood collection, rats were scored on the duration of time spent struggling during blood collection (s) (defined as rat moving in handler’s hands), number of vocalizations from video recordings, and number of times urinating and defecating during blood collection.

#### Statistical Analyses

Data were analyzed using RStudio (2020; Vienna Austria. URL: https://R-project.org/). For all response variables, normality was assessed using visual inspection of a histogram and quantile-quantile plot, as well as using the Shapiro-wilk test. Residuals of the models were also assessed using quantile-quantile plots and scatter plots. The experimental unit was cage.

##### Body weight

Data were normally distributed and not transformed. Effect of housing on rat body weight was assessed using generalized linear mixed models. Separate models were fitted for each time period weighed (day 0, 7, 14). Weight was included as the response variable, with fixed effects of treatment and sex, and random variable of cage.

##### HAT

Latency to touch observer and duration of touching observer were log_10_+1 transformed. Effect of housing and blood collection on response towards humans was assessed using generalized linear mixed models. Separate models were fitted for the response variables latency to touch and duration of interaction. The fixed effects were treatment and sex, and random variable was cage.

##### EPM

Data were normally distributed and not transformed. Effect of housing treatment was assessed using generalized linear mixed models. Separate models were fitted for response variables of anxiety-like behavior and locomotory activity. The fixed effects were sex and treatment, and random variable was cage.

##### Blood glucose

Glucose levels were log_10_1 transformed. Effect of housing environment on blood glucose levels was assessed using generalized linear mixed models, with glucose levels as the response variable, and fixed effects of treatment and sex, and cage as the random variable.

##### Behavior during blood collection

Number of vocalizations, number of urination and defecation events, duration of time struggling, and total duration of time to collect blood were measured. Total duration of blood collection was square root transformed. Duration of time struggling with log_10_+1 transformed. These measures were fitted in separate generalized linear mixed models with fixed effects of sex and treatment, and random variable of cage. Number of vocalizations and number of urination and defecation events during blood collection was low overall, therefore these data were analyzed using Poisson regression, with separate models fitted for vocalizations and for urination/defecation as the response variable, and treatment and sex as the fixed effects.

##### Behavior

Behavior, posture, and location of animals collected using scan sampling at the top of each hour was summarized using descriptive statistics. Behavior and posture collected during the active period (5:00) was used for statistical comparison between the housing groups. The following behavioral variables were square-root transformed for analysis: inactive, locomotion, grooming, social, cage exploration, vertical posture, lying posture. The following behavioral variables were log_10_+1 transformed: ingestion, aggression, sitting posture. Resource exploration and horizontal posture were not transformed. Abnormal behavior was not observed and was not included in the analysis. The effect of housing on behavior and posture was assessed using generalized linear mixed models. Separate models were fitted for each behavior and posture. Fixed effects included treatment, time period, and sex. Random effect was cage. Location is presented as proportion of time spent in each designated area of the environment. Social proximity data was analysed with binomial generalized linear mixed models due to the data being proportions of time near cage mates, with treatment, time period, and sex as fixed effects, and cage as the random effect.

### Study B

#### Animals and Housing

A total of 48 Crl:CD(SD) Sprague Dawley rats were used (24 male, 24 female) and animals were obtained from and housed at a pre-clinical safety assessment facility in Senneville, QC, Canada from May through August 2022. Animals were 54-61 days (∼7-8 wk of age) old at the start of the study. The duration of the study was 90 days. Animals were weighed weekly throughout the study and food, water, and environmental conditions were the same as for Study A.

Rats were housed in groups of 12 across 4 modified large animal cages (i.e., 2 enclosures each of males and females) as described for study A. The only modifications for study B were the addition of fleece (VetBed Canada, Smithers, BC) on the hammock and adding free-standing shelves high on the backs of the cages. On week 4 of Study B, 2 additional shelves were added (40cm L x 16 cm W, with a 2.5 cm ledge; 1,280 cm^2^ total area; Fig 4) to provide a total of 11,696 cm^2^, or approximately 974.67 cm^2^ per rat. Aside from the shelves, rats received the same in-cage resources as described in Study A.

**Fig 4.**
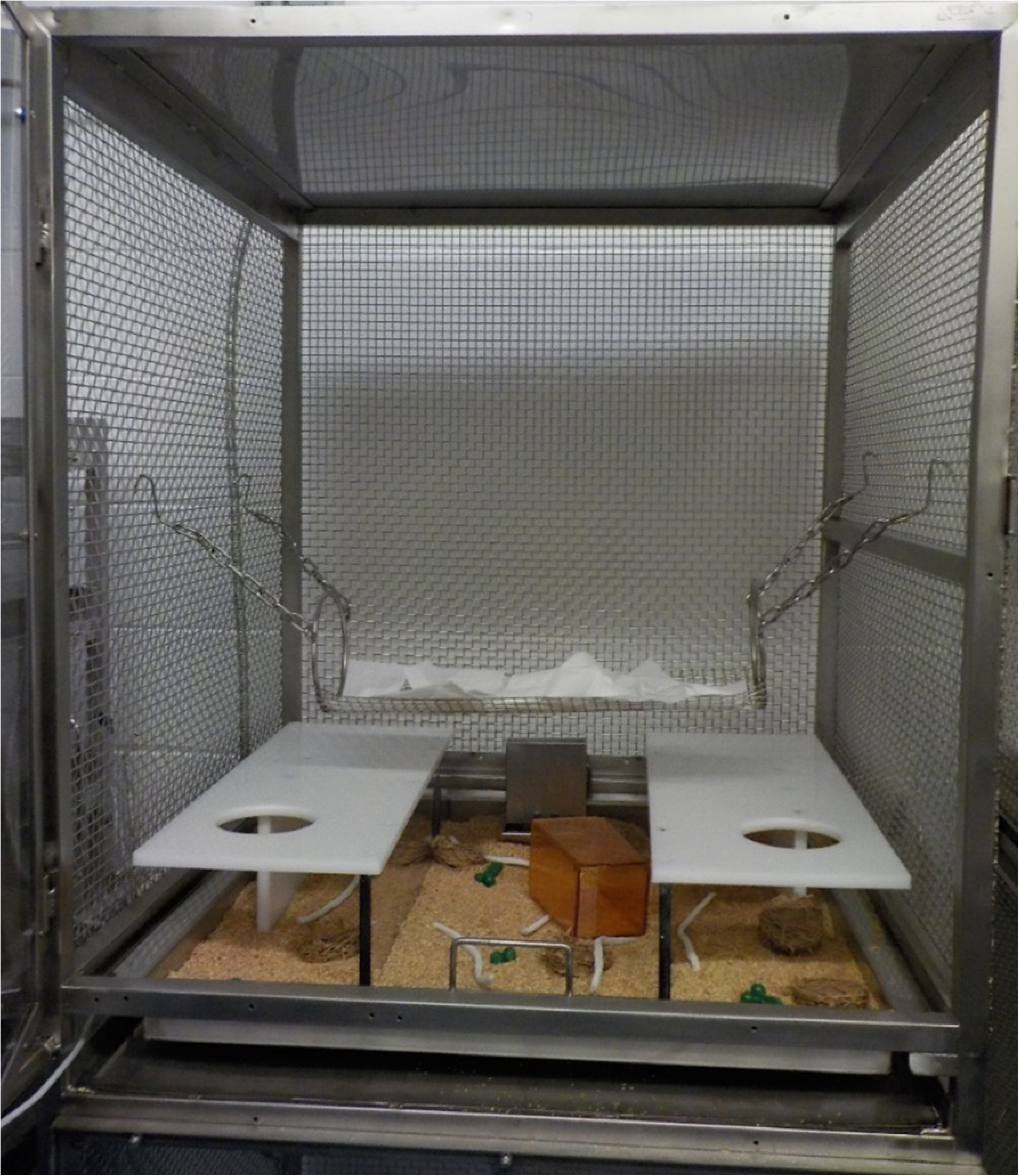
Image of the modified housing in study B with the addition of shelves on the back walls.

The control animals used for this study were selected from existing colony animals at the site who had received minimal handling. These animals remained in standard housing and were used for comparison in the behavior tests. There were 12 rats (6 male and 6 female) total across 4 cages (housed by sex with 3 rats per cage). Rats were 44 days (∼6 wk of age) of age at the start of the study. Standard housing for research rats included polycarbonate cages (24.1 cm L x 45.7 cm W x 20.3 cm H; 733.9 cm^2^) containing hardwood bedding. The cage contained 1 tunnel (Bio-Serv Rats Retreats^TM^, Certified, Flemington, NJ, USA), 1 half Nylabone© (Neptune City, NJ, USA) per rat, a rotating resource item (1^st^ week: Envigo Teklad 7979C.CS Certified/Irradiated Diamond Twists, Indianapolis, IN, USA; 2^nd^ week: timothy hay, Bio-Serv Certified, Flemington, NJ, USA), one 4 g nest pucks per rat (Bed-r’Nest puck, The Andersons, Delphi, IN, USA)

#### Gentle Handling

Rats were handled using the same procedure described in Study A except that there were 2 female handlers. The gentle handling procedure was the same as in Study A but with more rats per session in the transport box due to the increased number of rats per cage. Rats were collected 6 at a time from the modified cages using a spare standard rat cage (i.e., transport box). This procedure was followed until all rats in the transport box were handled. The remaining 6 rats were then brought into the transport box, and the same procedure was followed until all rats were handled, then repeated for each cage. The order of handling was selected at random at the cage level.

#### Rat Exercise Pen

Rats were given access to an exercise pen for 10-20 min once a week over 12 weeks during cage change of the modified primate cages. All 12 rats from the same cage were put into the play pen. The exercise pen was 157.5 cm L x 81.3 cm W, and was 78.7 cm off the floor. The pen contained (Fig 5) 1.36 kg of sani-chip (3 cm deep), 2 ladders, 4 tunnels (Bio-Serv Rat Tunnel Stainless Steel End Caps, Certified, Flemington, NJ, USA), 2 balls (Bio-Serv Jingle Balls^TM^, Certified, Flemington, NJ, USA), 2 pieces of 12 inch striped Maple wood saplings (Bio-Serv, Flemington, NJ, USA), 4 small wood gnawing blocks (Bio-Serv, Certified, Flemington, NJ, USA), 4 paper huts (Bio-Huts^TM,^ Certified Bio-Serv, Flemington, NJ, USA) and 12 pieces of tissue (Bio-Serv Rodent Nesting Sheets^TM^, Certified, Flemington, NJ, USA).

**Fig 5.**
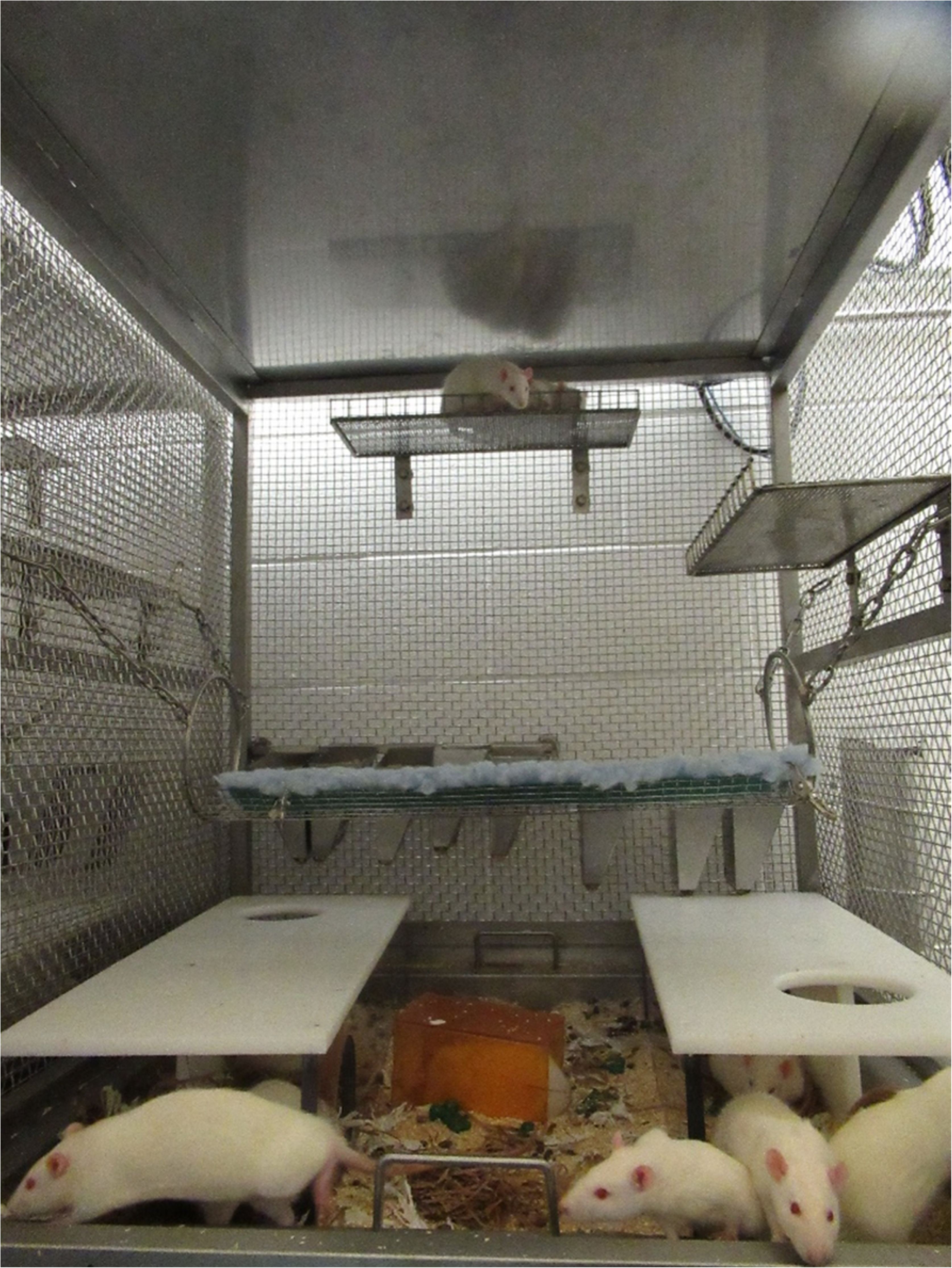
Rat exercise pen (157.5 cm L x 81.3cm W) used once weekly during cage changes for 10-20 min. The pen contained 1.36 kg of sani chip substrate, 2 ladders, 4 tunnels, 2 balls, 2 wood pieces, 4 wood gnawing blocks, 4 paper huts, and 12 pieces of tissue paper.

#### Behavior Tests

An elevated plus maze (EPM) test was used on day 86 to assess anxiety levels of rats as described for Study A.

A novel human approach test (HAT) was used to assess the rats’ relationship with humans as described for Study A. The HAT was conducted twice on day 89 of the study, in the morning and then again in the afternoon following the blood collection. The order of testing was randomized at the cage level then for individual in the cage, randomizing the animal numbers and taking the first 5-6 as one group, and the second 5-6 in the second group (depending on total number of animals in the cage). The same groups were tested together again after blood collection, but the order was randomized.

#### General Condition

Body weight (g) was collected individually on the day prior to the start of the study, then every 7 days throughout the study, ending on day 90. For the control rats, body weights were collected approximately every 2 wk. For comparison between the groups, weights were matched by age. Morbidity and mortality were also recorded throughout the study.

#### Blood Collection

Blood collection occurred on day 89 and individual rats were picked up by the handler who then sat in a chair with the fleece on their lap. The handler restrained the rat and collected blood from the saphenous vein. The rats’ rear legs were shaved for the procedure in the morning with blood collection occurring in the afternoon. A drop of blood was allowed to fall directly on a glucometer test strip as described for Study A. Rats were also scored on their struggle behavior during blood collection, vocalizations, and frequency of urination and defecation.

#### Behavior Scoring

The recording and scoring equipment used was the same as described for Study A. Order of videos watched, blinding, and intraobserver reliability were as described for Study A unless otherwise noted. There were 3 total observers and only one observer was responsible for each behavior measure.

Rat behavior was continuously recorded in the home cages for 24 hours one day per week during the study period for a total of 13 recordings (288 h). Behavior in the home cage was scored using the ethogram in Table 1 with instantaneous scan sampling every hour on the hour for the 24 h of the video. Rats were scored on overall behavior, location in the cage, and posture. These data from the first 4 weeks of observation were used to identify the rats’ most active hour (6:00am). During the active period, behavior was scored using instantaneous scan sampling every 30 s for the first 10 min of the hour, giving 20 observations per rat per week over 13 weeks. These data were summarized per week based on number of observations of that behavior/20. Behavior data were binned by time period, with period 1 (P1) spanning weeks 1-4, period 2 (P2) spanning weeks 5-8, and period 3 (P3) spanning weeks 9-13. The ethogram was the same as for Study A (Table 1), except the behavior climbing was added, defined as rat is on the walls of the cage. Postures and locations scored were the same as described for Study A except for in Study B, the location “Walls” included the horizontal shelves when they were added in week 4, and “Out of Sight” was added due to the number of animals and difficulty in confirming location in some instances. For study B, social proximity was scored the same as described for Study A. For all home-cage behaviors, there was a single observer. Confirmation of social proximity was done by a single, separate observer.

Rats were also scored for agonistic behavior using all occurrence sampling for total duration (s). To determine the optimal sampling strategy for recording agonistic behaviors, 100 videos were scored. Total duration of agonistic behaviors was summed per hour. The hours with the most agonistic behaviors were 4:00-7:00 and 18:00-22:00. These times were then scored for the remaining videos for total duration of agonistic behaviors (defined in Table 1).

The elevated plus maze (EPM) and human approach tests (HAT) were scored as described for Study A. The intraobserver reliability for the HAT was assessed by scoring 8 videos twice and comparing the results. The percentage of agreement for all observations combined was 85.82% (range from 80.19-93.80%).

#### Statistical Analyses

Data were analyzed using RStudio (2020; Vienna Austria. URL: https://R-project.org/). To compare test results between housing groups, Mann Whitney U tests were used due to the small sample size.

To compare behavior and posture across time period (3), Gaussian linear mixed models were used with sex and time period as fixed effects, and cage as the random effect. For all response variables, normality was assessed using visual inspection of a histogram and quantile-quantile plot. All behavior and postures (except horizontal posture) were not normally distributed and were square root transformed.

Normality of model residuals were checked using visual inspection of quantile-quantile plots. Least square means were compared across periods. P-values were adjusted for multiple comparisons using a Bonferroni correction. Location is presented as proportion of time spent in each designated area of the environment. Social proximity was analyzed as described for Study A, with sex and time period as fixed effects and cage as the random effect.

## Results

### Study A

#### General Condition

Throughout the study, there were no weight differences between rats in different housing treatments (day 0: F_(1,3)_ = 0.050, estimate = -1.241, SE = 5.529, *P*=0.826; day 7: F_(1,3)_ = 1.916, estimate= -11.475, SE = 8.289, *P*=0.190; day 14: F_(1,3)_ = 3.283, estimate = -15.442, SE = 8.522, *P*=0.094). On days 7 and 14, males were heavier than females (day 7: F_(1,3)_ = 13.199, estimate = 30.103, SE = 8.283, *P*=0.003; day 14: F_(1,3)_ = 38.733, estimate = 52.932, SE = 8.494, *P*<0.0001). There was no sex effect on day 0 (*P*=0.092). No mortality or morbidity was reported during the study period for either housing type.

#### Response to Humans

There was no effect of housing (*P*>0.501) or sex (*P*>0.856) on latency to approach the human before (F_(1,3)_ = 0.002, estimate = 0.005, SE = 0.111, *P*=0.966) or after blood collection (F_(1,3)_ = 0.4555, estimate = 0.091, SE = 0.134, *P*=0.501) during the human approach test (HAT), but rats in both housing groups were quicker to approach before blood collection than after (F_(1,4)_ = 7.025, estimate = -0.215, SE = 0.081, *P*=0.009). Rats also spent less time interacting with the human before blood collection than after (F_(1,4)_ = 4.977, estimate = -0.109, SE = 0.049, *P*=0.028). T rats spent less time interacting with the human in both HATs compared with C rats (F_(1,4)_ = 7.143, estimate = -0.139, SE = 0.052, *P*=0.009). Male rats from both groups spent less time interacting with the human than females (F_(1,4)_ = 14.542, estimate = - 0.194, SE = 0.050, *P*=0.0002).

There were no differences in blood glucose levels between sexes (*P* = 0.130) or housing treatments (*P*=0.166). There were no differences in struggle behavior during blood collection based on sex (*P*=0.435) or housing treatment (*P*=0.801). There was an effect of housing treatment on number of urination and defecation events and number of vocalizations during blood collection. T rats had more urination and defecation events (F_(1,2)_= 7.929, estimate = 2.356, SE = 1.026, *P*=0.022) and vocalized more compared to C rats (F_(1,2)_ = 8.707, estimate = 0.465, SE = 0.164, *P*=0.005).

#### Elevated Plus Maze (EPM)

There was an effect of housing on anxiety-like behavior during the EPM (Fig 6). T rats spent more time in the open arms of the EPM compared to C rats (F_(1,3)_ = 5.287, estimate = 0.139, SE = 0.061, *P*=0.038). There was a sex effect, with males spending less time in the open arms of the EPM than females (F_(1,3)_ = 15.464, estimate = -0.238, SE = 0.060, *P*=0.002). There was no effect of housing on locomotor activity (*P*=0.133) but there was a sex effect, with male rats showing less locomotor activity compared to female rats (F_(1,3)_ = 7.689, estimate = -3.349, SE = 1.197, *P*=0.016).

**Fig 6.**
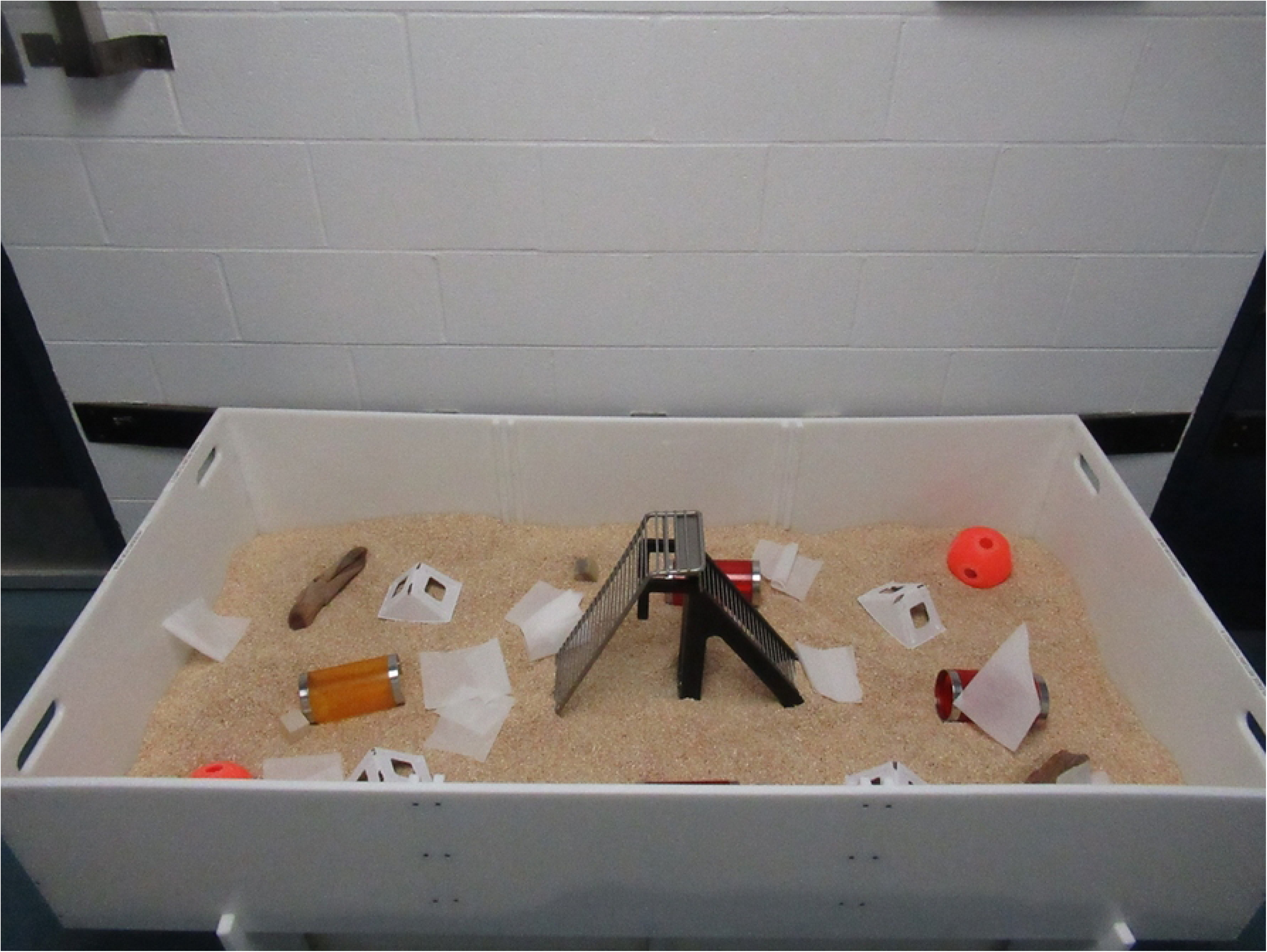
Anxiety-like behavior displayed in rats in standard housing (C) or in modified primate cages (T). Anxiety-like behavior was assessed using an elevated plus maze, where duration of time spent in the open arms of the maze compared to total amount of time in all arms of the maze is indicative anxiety levels. Data are presented as raw means and standard errors.

#### Overall Behavior

The overall behavior of the rats summarized over the entire study period by housing treatment is presented in Fig 7. T rats spent less time inactive (F_(1,4)_ = 31.865, estimate = -3.569, SE = 0.632, *P*<0.0001), consuming food (F_(1,4)_ = 5.568, estimate = -0.195, SE = 0.083, *P*=0.038), grooming (F_(1,4)_ = 23.761, estimate = -3.632, SE = 0.745, *P*=0.0003) and less time on social behavior (F_(1,4)_ = 15.899, estimate = -1.984, SE = 0.498, *P*=0.002) and cage exploration (F_(1,4)_ = 15.278, estimate = -1.586, SE = 0.406, *P*=0.002) compared with C rats. T rats spent more time on resource exploration (F_(1,4)_ = 21.433, estimate = 59.893, SE = 12.928, *P*=0.0003) and locomotion (F_(1,4)_ = 171.002, estimate = 8.868, SE = 0.678, *P*<0.0001) compared with C rats. Insufficient agonistic behavior was recorded to be further analyzed (mean: C = 0.18, T = 0.1).

**Fig 7.**
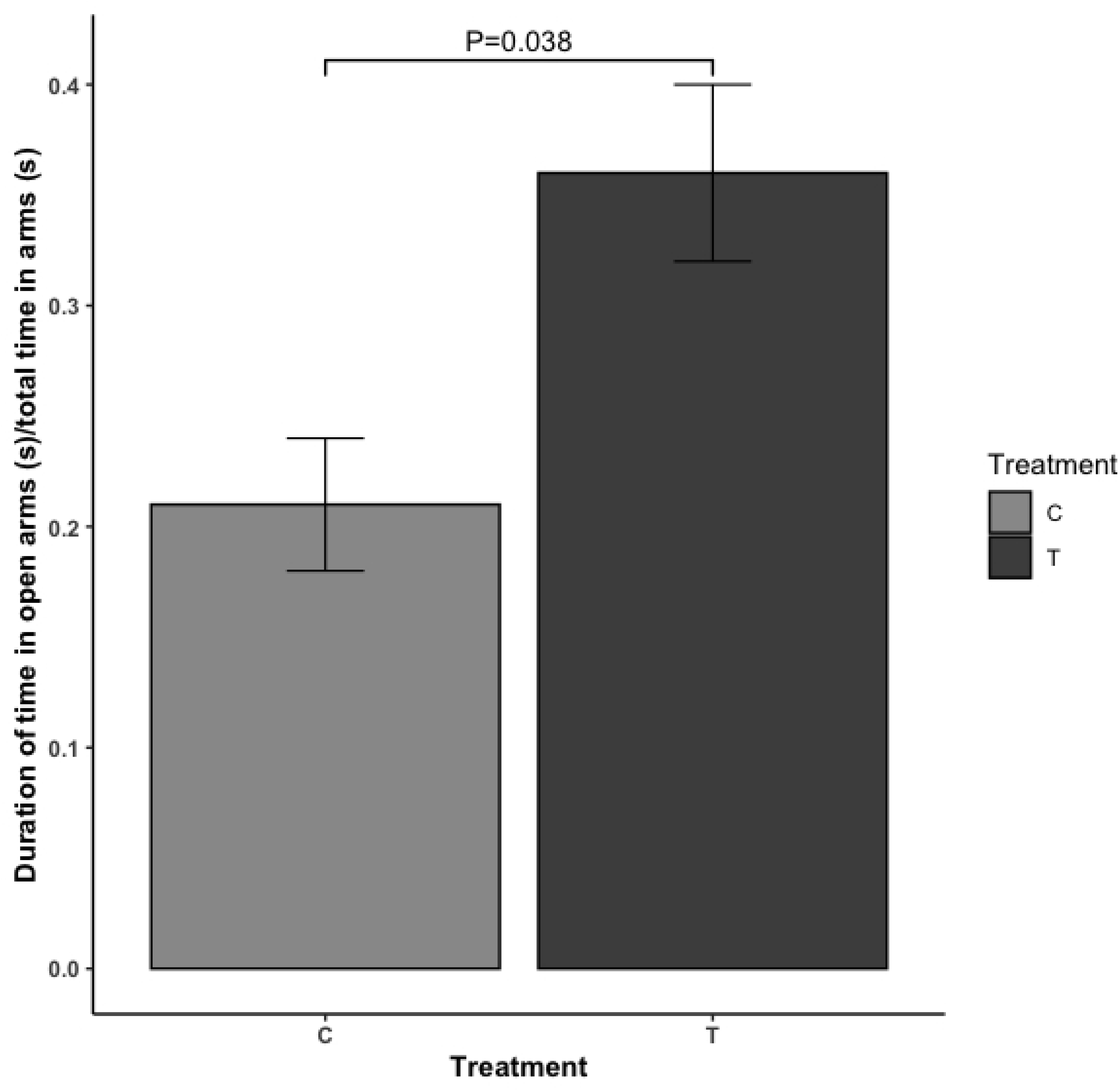
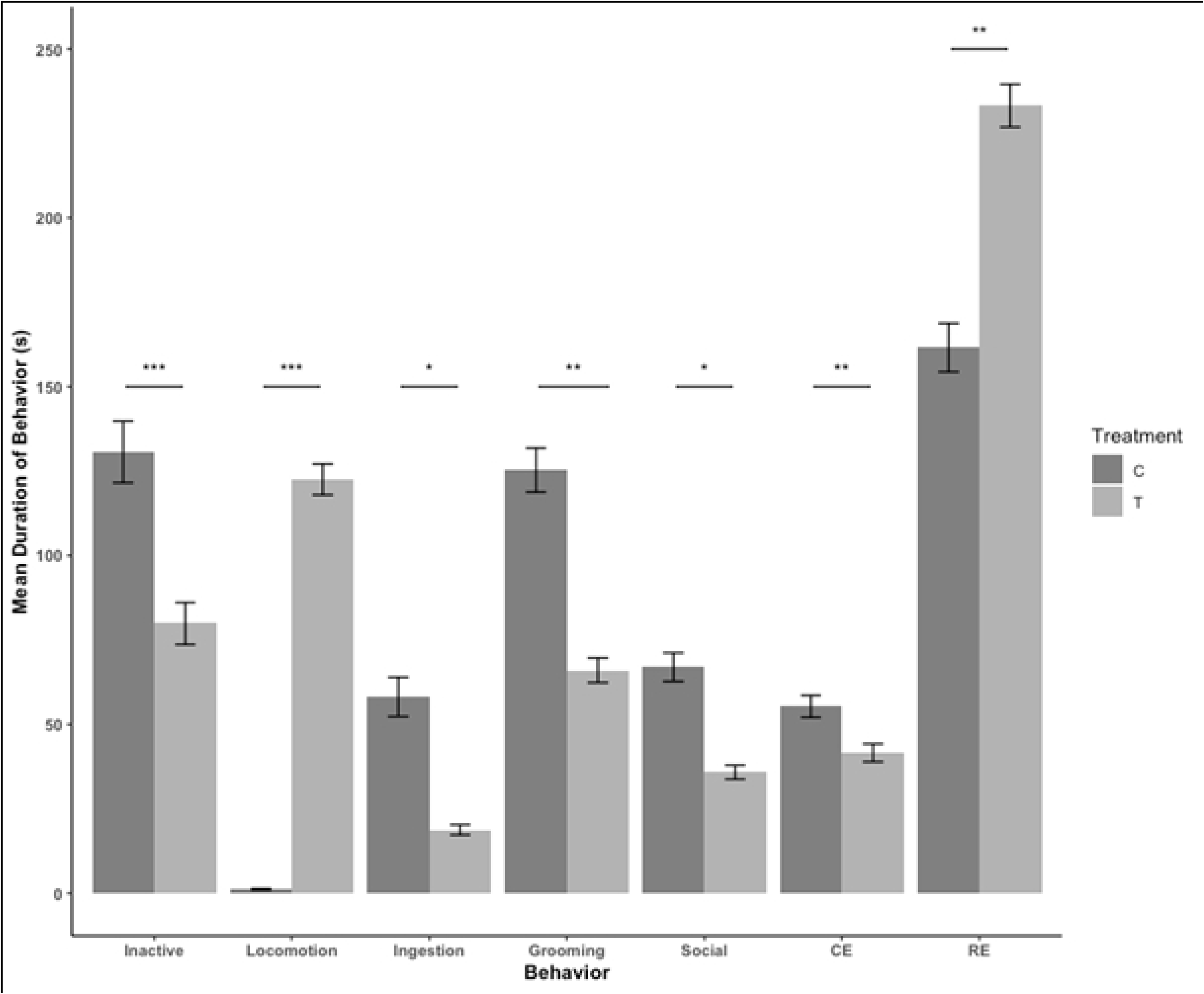
Overall summary of rat behavior in study A, comparing behavior of rats in standard housing (C) and rats in modified primate cages (T). CE: cage exploration, RE: resource exploration. Data are presented as raw means and standard errors. Behavior was scored using duration of time spent on each behavior during the first 10 min of the active period (5:00) for each day of the 18-day study period, then summarized for the entire study period and compared between housing groups using generalized linear mixed models. *=P<0.05, **=P<0.01, ***=P<0.0001

Sex differences were found in that male rats were more inactive (F_(1,4)_ = 19.595, estimate = 2.762, SE = 0.620, *P*=0.0008) and spent less time on cage exploration (F_(1,4)_ = 20.127, estimate = -1.805, SE= 0.401, *P*=0.0007) compared with female rats. There was also an effect of time period. During period 2, rats in both housing treatments spent less time moving (F_(1,4)_ = 27.049, estimate = -1.510, SE = 0.290, *P*<0.001) and exploring resources (F_(1,4)_ = 51.451, estimate = -61.497, SE = 8.569, *P*<0.0001), while spending more time grooming (F_(1,4)_ = 3.889, estimate = 0.679, SE = 0.344, *P*=0.049), social behaviors (F_(1,4)_ = 61.917, estimate = 2.233, SE = 0.284, *P*<0.0001), and exploring their cage (F_(1,4)_ = 21.290, estimate = 1.235, SE = 0.268, *P*<0.0001) compared to period 1.

#### Posture

Differences were observed in posture between C and T rats. There were no differences in vertical posture (*P*=0.167). T rats spent more time in horizontal posture (F_(1,4)_ = 32.751, estimate = 113.950, SE = 19.890, *P*<0.0001), but less time in lying (F_(1,4)_ = 70.068, estimate = -5.873, SE = 0.701, *P*<0.0001) and sitting (F_(1,4)_ = 20.879, estimate = -0.631, SE = 0.138, *P*=0.0006) postures compared with C rats. There was a sex effect for vertical and lying postures, with males spending less time in vertical posture (F_(1,4)_ = 6.457, estimate = -1.613, SE = 0.634, *P*=0.024) and more time in lying posture (F_(1,4)_ = 21.705, estimate = 3.238, SE = 0.692, *P*=0.0005) compared with females. There also was an effect of time period for vertical and lying posture, with rats in general spending less time in vertical posture (F_(1,4)_ = 8.191, estimate = - 0.791, SE = 0.276, *P*=0.004) and more time in lying posture (F_(1,4)_ = 8.021, estimate = 1.316, SE = 0.465, *P*=0.005) in period 2 compared with period 1.

C rats spent on average 65% (SD=29) of observations in proximity of cage mates, while T rats spent on average 80% (SD=20) of observations in proximity to cage mates. There was no statistical difference in social proximity between treatment groups (*P*=0.293). There was a sex effect, with females spending more time in proximity of cage mates, compared with males (*P*<0.0001), and generally, rats spent less time in close proximity of cage mates in period 2 (*P*=0.038).

#### Location

Time spent in each location of the cage were also summarized across study day for C rats (Fig 8a) and T rats (Fig 8b). C rats spent most of their time in the back of the cage. T rats spent most of their time under the left and right tunnelss, and on the floor of the cage.

**Fig 8.**
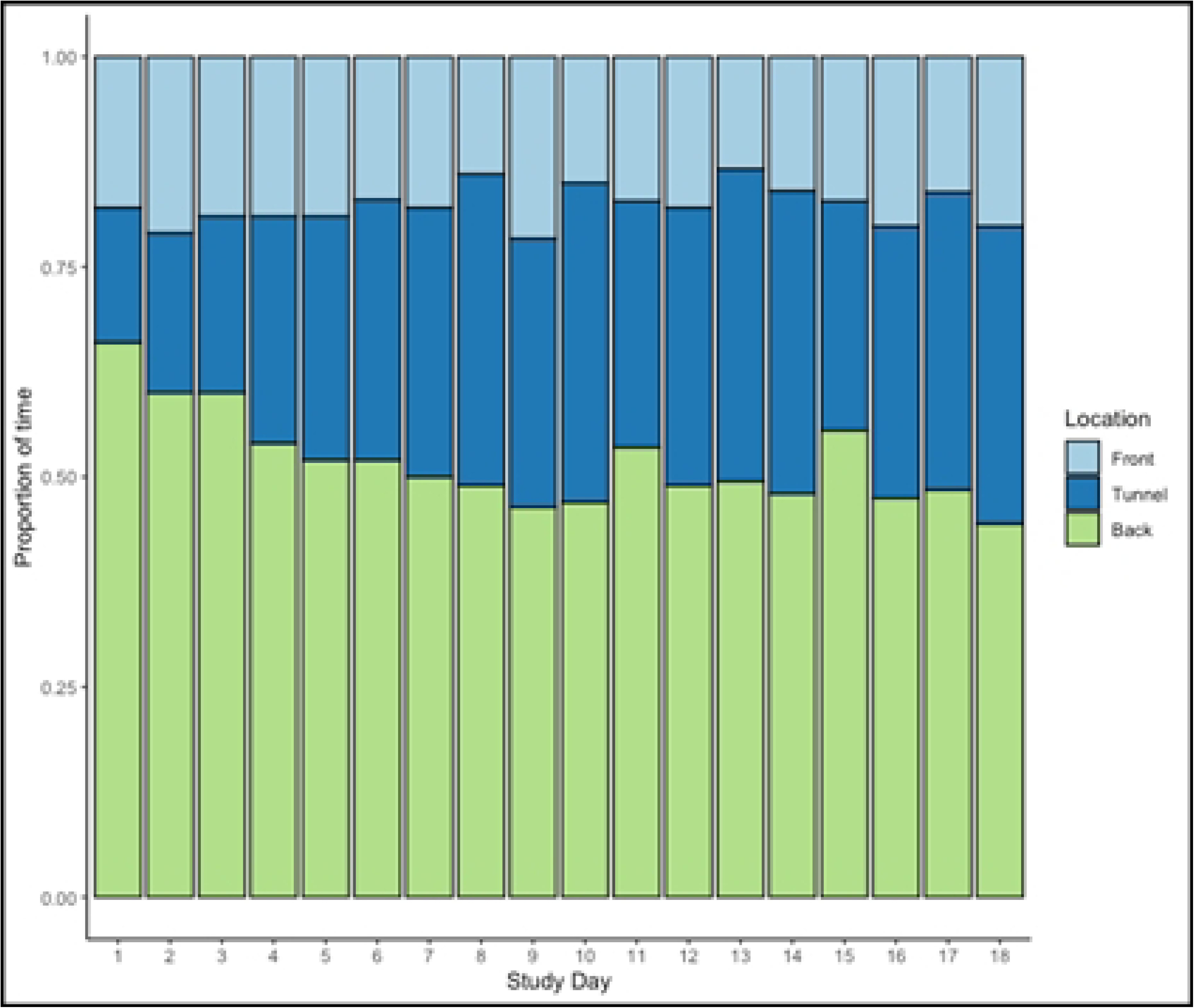
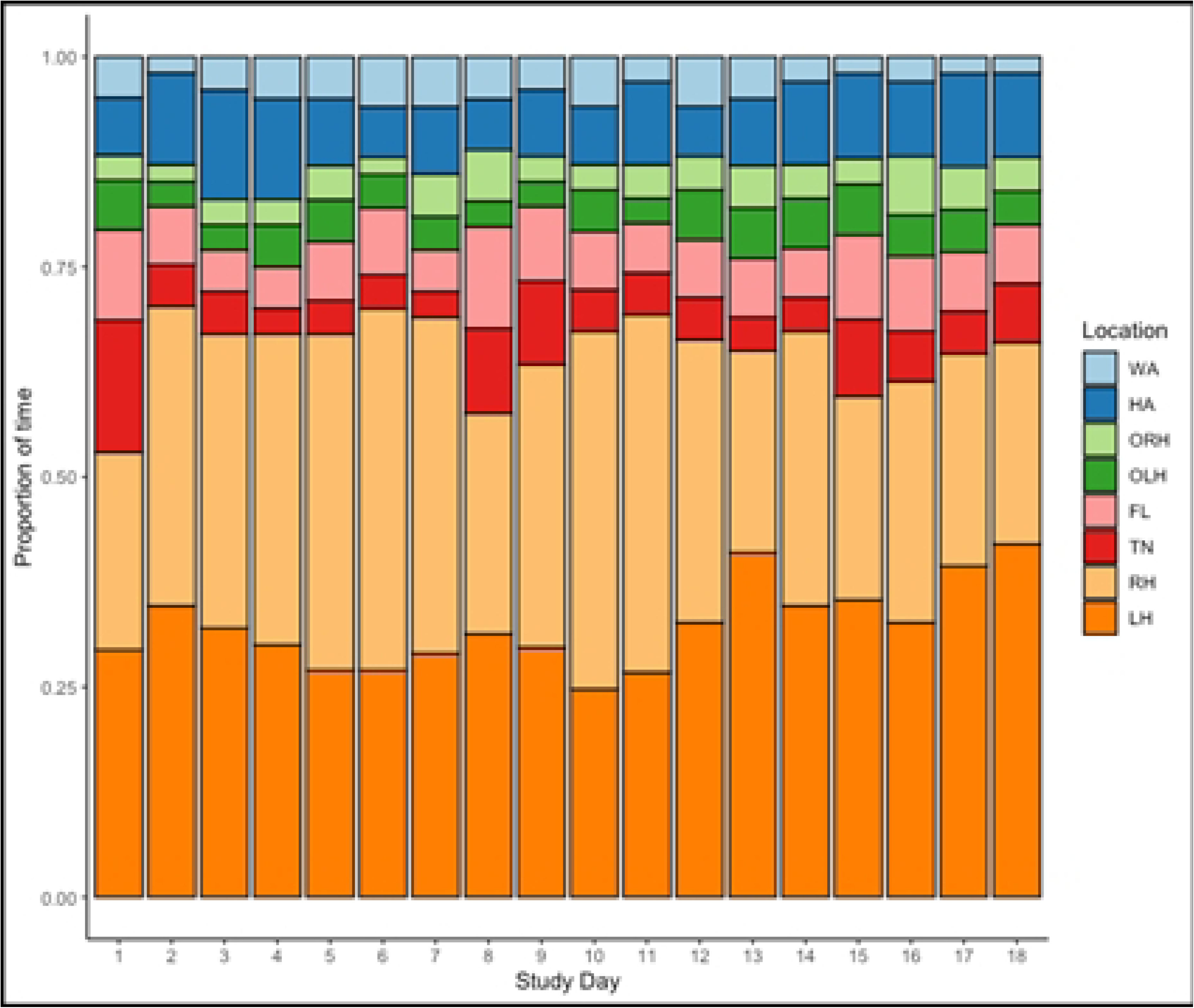
Time spent in each location of the cage for rats in standard housing (C, 8a) and rats in modified primate cages (T, 8b). For T rats, the locations include LH: under left hut, RH: under right hut, TN: in tunnel, FL: floor, OLH: on left hut, ORH: on right hut, HA: hammock, WA: on walls. Location was scored using scan sampling recording number of animals in each location at the top of the hour every day for the study period, then summarized by day. Data are presented as proportion of time spent in each location.

### Study B

#### General condition

No difference in weight was seen between the control rats housed in standard housing and those in the modified primate cages over the 90-day study period (W = 270, P=0.829). At the end of the study, the male rats in the T housing weighed 663.7 g (SE = 10.4) and females weighed 403.3 g (SE = 7.5) on average.

There were 8 reports of morbidity, which all occurred in male cages between study weeks 3-7. The conditions were skin lesions, possibly indicative of fight wounds (6 occurrences). There were 2 reports of injured toes and limping observed in week 6 in one male cage. This was presumed to be related to the elevated shelves. The shelves were adjusted and lowered, and no further toe or limb injuries were observed. Two incidences of mortality were reported for male rats, one in week 9 and one the final day of the study in week 13. One mortality occurred immediately after a handling event. The second incident occurred on the last day, where the rat was found under the left hut of the cage. Necropsy revealed no gross findings in either case and there were no observations to explain the mortalities.

#### Response to Humans

There was no difference in blood glucose levels between rats in different housing treatments as a result of restraint (*P*=0.801).

Rats in the modified housing had a longer latency to approach the person (W = 737, *P*=0.012) and spent less time in contact with the person compared with control rats (W = 1730, *P*<0.0001) in the HAT. There was no difference in HAT latency (control: *P*=0.839; modified cages: *P*=0.614) or touch duration (control: *P*=0.291, modified cages: *P*=0.568) between the HAT before blood collection and the HAT after, showing no effect of restraint for blood collection (Fig 9).

**Fig 9.**
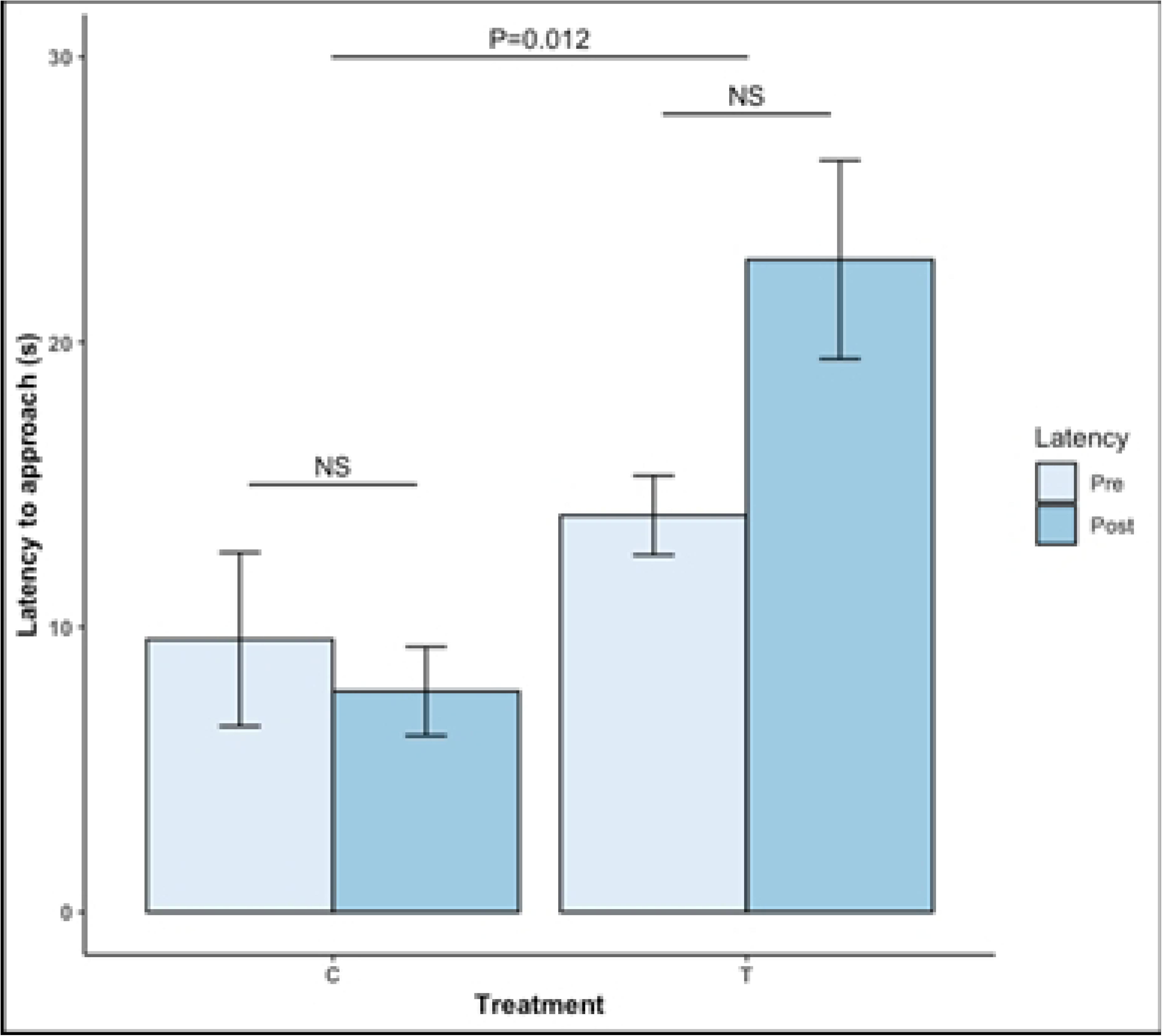
Rat responses during the human approach test, comparing rats housed in standard housing (C) and rats in modified large animal cages (T). Rats were tested for 1 min prior to blood collection (pre) and after blood collection (post) by measuring their latency to approach the human handler (9a) and duration of time in contact (9b). Data are presented as raw means and standard errors.

#### Elevated Plus Maze (EPM)

Rats in the modified housing showed less anxiety by spending more time in the open arms of the maze compared to control rats (control: mean = 0.08, SE = 0.01; modified: mean = 0.38, SE = 0.03; W = 40, P<0.0001), but there was no difference in locomotion between groups (W = 268.5, P=0.984).

#### Overall Behavior

Rats in the modified cages did show changes in behavior over time (Fig 10), becoming more inactive with each period and spending less time on other behaviors such as locomotion, eating, grooming, social behaviors, cage exploration, and resource exploration. There was an increase in climbing behavior between P1 and P2, then climbing remained stable for the remainder of the study.

**Fig 10.**
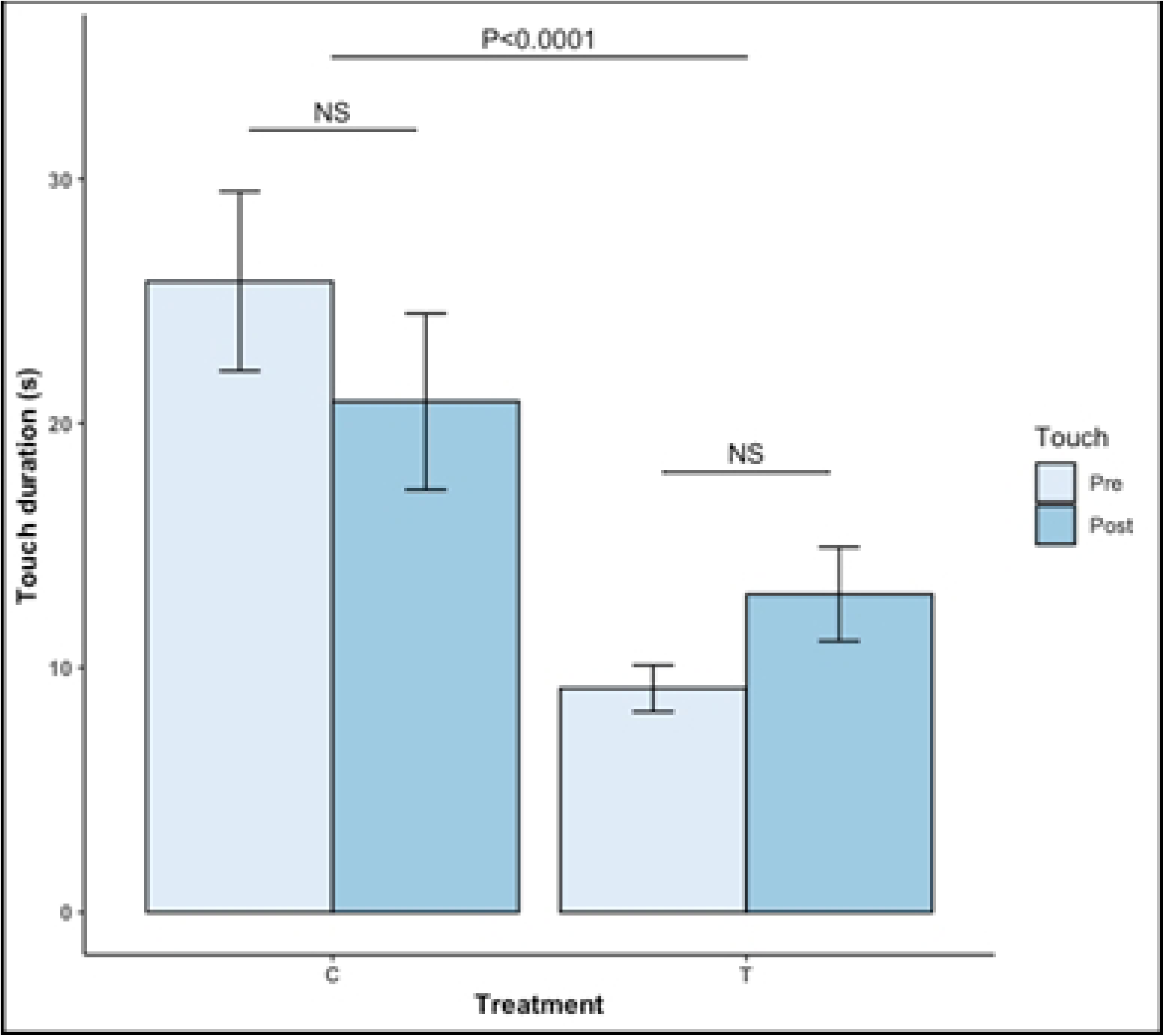
Overall summary of rat behavior in study B, showing how behavior changes over 3 time periods of 30 days each (CE: cage exploration, RE: resource exploration). Data are presented as raw means and standard errors. Behavior was scored using duration of time spent on each behavior during the first 10 min of the active period (6:00) one day per week over 90 days, then compared across time periods using generalized linear mixed models. *=P<0.05, **=P<0.01, ***=P<0.0001

Rats on average spent 84% (SD=13) of their time in proximity with cage mates. There were no differences based on sex (*P*=0.917) or time period (*P*>0.557).

#### Agonistic behavior

Agonistic behavior was observed throughout the study, being highest in P1 (P1: mean = 261.71, SE = 30.66; P2: mean = 176.71, SE = 23.90; P3: mean = 134.93, SE = 17.06). There was a significant decrease in this behavior between P1and P2 (estimate = 3.26, SE = 1.04, *P=*0.005) and P1 and P3 (estimate = 5.16, SE = 0.988, *P*<0.0001), and it remained stable between P2 and P3 (estimate = 1.01, SE = 0.969, *P*=0.121). There were sex differences, with female rats displaying more agobistic behavior (estimate = -5.165, F_(1,4)_ = 40.656, *P*=0.024).

#### Posture

Similar to the behavior patterns, rats spent less time in active postures over the course of the study and more time in resting postures. Vertical posture decreased between P1 and P2 (estimate = 0.089, SE = 0.016, *P*<0.0001) and P1 and P3 (estimate = 0.102, SE = 0.015, *P*<0.0001) but remained the same between P2 and P3 (estimate = 0.014, SE = 0.015, *P*=0.614). Time spent in horizontal posture decreased for each time period (P1-P2: estimate = 0.147, SE = 0.032, *P*<0.0001; P1-P3: estimate = 0.332, SE =0.031, P<0.0001; P2-P3: estimate = 0.185, SE = 0.030, *P*<0.0001). Lying postures increased across each time period (P1-P2: estimate = -0.155, SE = 0.028, *P*<0.0001; P1-P3: estimate = -0.322, SE = 0.027, *P*<0.0001; P2-P3: estimate = -0.167, SE = 0.026, *P*<0.0001). Time spent sitting increased between period 1 and 2 but stayed the same between period 2 and 3 (P1-P2: estimate = -0.046, SE = 0.017, P=0.015; P1-P3: estimate = -0.042, SE = 0.016, *P*=0.024; P2-P3: estimate = 0.005, SE = 0.016, *P*=0.949).

#### Location

Rats initially spent most of their time under or on the huts (54% of time on or under left or right hut in P1; P2: 38%; P3: 30%), but then began spending their time on the shelves after they were added (P1: 8%; P2: 26%; P3: 38%) (Fig 11).

**Fig 11.** Time spent in each location of the cage for rats in Study B. Locations include LH: under left hut, RH: under right hut, TN: tube/tunnel, FL: floor, OL: on left hut, OR: on right hut, HA: hammock, WS: walls (including shelves), OS: out of sight. Location was scored using scan sampling recording number of animals in each location at the top of the hour once a week over 90 days, then summarized by each 30-day time period (3 in total). Data are presented as proportion of time spent in each location.

## Discussion

This research aimed to explore the use and impact of modified primate cages as rat housing in 2 studies. In study A, behavioral and physiological parameters between rats housed in different housing environments over 18 days were compared. Rats in the modified primate housing had reduced anxiety and spent more time on active behaviors. There were no differences in latency to approach a human based on housing method, but rats in the modified primate housing spent less time in contact with the person during the HAT test and displayed more behavioral stress indicators during blood collection. Over the 18-day study period, the rats in the modified primate cages became less active over time, suggesting that rats may habituate to the housing. Thus, a follow up study was conducted (Study B) to observe rats in the modified primate cages over 90 days to evaluate safety as the animals became heavier and habituation or changes in behavior. Rats became more inactive over the 90-day study period, but continued using the resources provided, including daily climbs up and down to the elevated shelves, suggesting sustained benefits of the housing on rat welfare.

These results are comparable to other studies looking at housing environment, where rats provided with more in-cage resources are more active and exploratory compared to rats housed in resource-barren cages [27-29]. It has also been observed that stress decreases locomotion and exploration [30,31], suggesting that rats housed in complex environments express more natural behavior, which are beneficial for promoting positive affective states [32,33]. In the present study, rats housed in standard cages engaged in more cage exploration, self-grooming, and eating compared with rats in the modified housing, potentially indicative of boredom or stress [27,34,35]. In a study by Abou-Ismail and colleagues (2010), exploration was defined as sniffing the cage wall, cage top, and sniffing air outside the cage, as defined in the present study [27]. An increase in this type of exploration for rats housed in a barren environment, as well as increased bedding-directed behaviour, was observed, suggesting that rats in barren cages did not have enough stimulation, causing them to occupy themselves with basic items in the cage [27].

The CCAC housing guidelines for rats indicate that cage height should allow rats to stretch vertically, and therefore should accommodate the total body length of the rats plus 4 cm to account for the full stretching posture [11]. This is the first regulatory precedent for this standard of housing despite empirical evidence of the importance of standing vertically to rat natural body postures [4,36], research showing that cages with multiple levels of varying cage heights is beneficial for welfare [4,37-39], and knowledge that standard rat shoebox caging does not allow them to stand fully upright. In the present study, there were no difference between housing groups in time spent in vertical posture but there were other measures suggesting the modified primate housing was beneficial to rat welfare such as less time spent in lying and sitting postures, ess time inactive, and demonstration of less anxiety-like behaviors. A recent systematic review discusses the topic of boredom and its negative effects on rodent welfare [40].

The authors specifically describe awake-inactivity as a measure of boredom, which is similar to what was found in the current study regarding control animals spending more time lying, sitting, and inactive, which could have effects on rat health and model validity [40]. These results taken with other literature on the benefits of multi-level housing may suggest that the primary factor improving rat welfare may not be whether they can spend more time in a vertical posture, but instead in the choice and control offered by housing with additional space and resources [41,42]. In Study B, even though rats became more inactive over time, they continued to use the additional resources provided, in particular, the elevated resting areas, which became a preferred location after being introduced in the second study period. A study investigating age-related differences in emotional behavior in rats demonstrated that rats showed a decrease in motor activity with age but did not find any changes in social interaction with age [43]. The current study did observe a decrease in active social interaction as well as decreased agonistic behaviors with time. In natural conditions, rats display social dominance and fighting becomes less frequent when a hierarchy is established. This may explain the decrease in agonistic behavior from P1 to P2 with a dominance hierarchy established and the stable baseline agonistic behavior seen between P2 and P3 [44], although it should be noted that scuffles and fighting never resulted in the need to separate animals because of wounding severity, despite housing 12 animals per enclosure. The current study also demonstrated that females expressed more aggression than males, which is not typically seen in the literature. However, play scuffling could have been mistaken for agonistic behavior during video scoring and is seen at a higher rate in juveniles and females [44]. In a research setting, aggression needs to be monitored to ensure the safety and welfare especially in large groups, but it is important to acknowledge that large groups offer an ecologically valid environment in comparison to standard housing. Despite a decrease in active social behaviors, rats remained in close proximity of one another, choosing to spend time in the same locations. Rats in T housing spent 80% of their time in close proximity to cage mates, compared with 65% of time spent in proximity in the C group. Though not statistically significant, this shows that despite having more space to get away from cage mates, rats chose to be near conspecifics. Time spent in proximity to cage mates did not decrease over time for rats in the T treatment, showing continued motivation for social interaction as rats aged [43].

The physical state of the animal is a common animal-based measure used to assess health and welfare [11]. Standard housing has been linked with increased rodent obesity [6,45,46]. Two studies looked at the effects of the removal of resources in male and female rats and both demonstrated that body weight increased for weeks after the removal of in-cage resources due to the development of hyperphagia and other depression-like behaviours, dysregulation of the adrenocortical axis, and increased helplessness behavior [47,48]. This suggests that rats housed in an environment with increased space, resources, and conspecifics may have more lean muscle mass due to increased energy expenditure and reduced food intake. In this study, there were no weight differences between rats housed in standard cages versus the resource-enriched cages in either study A or B. Body composition was not evaluated and additional measures may be needed to understand the impact of more complex housing on physical condition of the animals, such as body composition analysis.

Mazhary and Hawkins (2019) collected opinions and attitudes regarding rat cage heights from laboratory animal personnel in the UK, and the major concerns expressed about implementing taller cages included handling and safety, with taller cages typically being heavier and more difficult to handle [49]. Changes to rack height could lead to personnel injuries when moving cages from the top or bottom shelves, financial concerns of replacing housing and racks, animal welfare concerns for rats receiving surgery or implantations, concerns that the animals would be more territorial and aggressive towards handlers, and human perceptions related to rat behavioral needs [49]. At the research facility where the present study was conducted, there were a large number of outdated stainless steel primate cages out of use used. The aim of repurposing these cages for rat housing was to improve ergonomic conditions compared to larger rat cages, repurpose old equipment to reduce waste, provide cost savings by utilizing existing resources, and overall improve rat welfare by providing additional space and resources to improve behavioral and physiological outcomes. The regular gentle handling protocol used in the study was also implemented to address concerns over human-rat interactions.

Despite receiving the same gentle handling protocol, there were differences in response to humans observed in Study A, with rats in the modified housing spending less time in contact with the person during the HAT and being more likely to vocalize, urinate, and defecate during blood collection. Similar results were seen in Study B, even with control rats not receiving the gentle handling, where rats in standard housing were quicker to approach and spent more time in contact with a person. This may not be indicative of a different association with humans as no bites or scratches occurred, but rather a difference in motivation [50,51]. Rats in the standard housing may be more willing to interact with people as a form of stimulation, whereas rats in modified housing may be less interested in human interactions due to behavioral and psychological needs being met by their home environment and the social interactions they experience there. It is also important to factor in the testing environment [33]. While latency to approach and time interacting are considered reliable indicators of the valence of human interactions for rats, it is important to note that willingness to interact may not always be a clear indicator of positive human association [50]. Standard housing for rats can increase anxiety and depressive-like states [40], therefore the lack of response during restraint for blood collection could indicate a freeze response or learned helplessness, rather than a lack of aversion to the procedure [52,53]. The display of fear behaviors during restraint may also indicate a greater willingness to express emotional states from animals with less anxiety and who have been habituated to people. A few studies have looked at cognitive bias in rats in relation to anxiety and environmental resources and it was found that rats exposed to a negative environment react less to an ambiguous stimulus in comparison to rats exposed to a calmer environment that react more to the ambiguous stimuli [54,55]. Barrata and colleagues (2023), explain theories on neurological pathways in relation to behavior testing in which rats that have more control exhibit increased resiliency when faced with adverse stimuli [52]. However, the relationship between housing, affective states, and human interactions and how these factors might influence each other are not well understood at this time. Future research should investigate these topics in more depth as well as develop measures of positive affective states.

Increasingly, the importance of providing resources to research rodents has been recognized from an ethical, animal health (physical and mental), and scientific perspective [6] and is being incorporated into regulatory requirements [11]. Increasing demands surrounding animal welfare can be met with negative opinions, that as the belief that this takes away initiative and personal responsibility [56] or differences in opinion regarding animal behavioral needs [49], particularly between different stakeholders [57]. There are other limitations to implementing new animal management practices such as the cost of the resources and labor, practicality, and scientific variation [6,57,58]. The lack of reporting on housing and resources used in rodent studies makes it difficult to assess impacts on cost and feasibility at this time but may be better in the future due to the updated ARRIVE guidelines [7,59]. In the present study, prior to study start, animal care technicians believed there would an increase in the time to change/clean modified cages compared with the standard cages (*personal communication,* August 2022). However, the larger enclosure and increased bedding did not get soiled as quickly compared to the standard cages, reducing cage change frequency. The modified cages were cleaned similarly to large animal cages, being sent to the cage wash every 2 weeks. In comparison, standard cages are required to be changed twice weekly, thus the time is similar. Another important factor is staff satisfaction in relation to animal welfare, where being able to provide additional resources and enrichment to animals can improve job satisfaction and reduce feelings of compassion fatigue [57,60-62]. In this study, researchers as well as site technicians and veterinarians reported that seeing the rats express natural behaviours such as climbing, vertical posture, and social interaction in a large group improved their job satisfaction suggesting that increase time to provide this environment benefits the moral feelings of using animals in research (*personal communications,* August 2022). Implementing new housing and husbandry practices in a research setting requires collaboration between many stakeholders including research personnel, regulatory experts, management, and animal support staff [56].

Providing larger, better resourced housing for research rats is an important step to improve their welfare. Future studies should explore the practicality of using this type of housing for rats on a research study. One limitation of this study was that the rats were not actively enrolled in a preclinical study. The addition of study procedures may impact rat behavior, particularly their response to humans, and therefore needs to be explored further. In Study B, there were 2 incidences of mortality that occurred. While there was no evidence that the mortality was caused by the housing, more long-term observations of rats in modified large animal cages could help further understand the full range of risks and benefits to rats.

## Conclusion

In conclusion, improved housing created by modifying primate cages for rats resulted in greater expression of natural behaviors, such as locomotion and resource exploration, and less anxiety compared with rats housed in standard polycarbonate cages. Rats in the modified housing also spent less time inactive, and in lying and sitting postures suggesting overall greater activity. Although rats in the modified housing did demonstrate reduced activity over time, they continued using the additional resources provided, suggesting sustained welfare benefits. There were differences in response towards people based on housing, whereas rats in the modified housing spent less time in contact with people during an HAT and showed more stress behaviors during blood collection, despite regular gentle handling. Despite this, there were no bites or scratches or evidence of aggression towards people. Overall, the modified primate cages provided more opportunities for expression of natural behaviors and postures, and had positive welfare benefits that should be considered in future decisions on research rat housing.

## Declaration of Interest

The authors work for or have worked for Charles River and declare that this study received funding from Charles River. The funder was not involved in the study design, collection, analysis, interpretation of data, the writing of this article or the decision to submit it for publication. The authors declare no conflict of interest.

## Acknowledgements

This work was presented at the ANZCAART Conference, 8 to 10 August 2023, Adelaide, AUS.

The authors would like to thank personnel at the Senneville facility for their assistance in completing these studies.

## Notes

### Competing Interest Statement

The authors have declared no competing interest.

## References

1. Russel WMS, Burch RL. The Principles of Humane Experimental Technique. London, UK: Methuen; Potters Bar, UK; Herts, UK: UFAW; 1959.

2. National Centre for the Replacement, Refinement, and Reduction of Animals in Research (NC3Rs). The 3Rs. https://nc3rs.org.uk/who-we-are/3rs. Accessed [March 26, 2024].

3. Turner PV. Moving Beyond the Absence of Pain and Distress: Focusing on Positive Animal Welfare. ILAR J. 2021;60(3):366–372. doi:10.1093/ilar/ilaa017.

4. Makowska IJ, Weary DM. The importance of burrowing, climbing and standing upright for laboratory rats. R Soc Open Sci. 2016;3:160136. doi:10.1098/rsos.160136.

5. Makowska IJ. The Rat. In: Sørensen DB, Cloutier S, Gaskill BN, editors. Animal-centric Care and Management: Enhancing Refinement in Biomedical Research. Boca Raton, FL: CRC Press; 2021. p. 121–134.

6. Cait J, Cait A, Scott RW, Winder CB, Mason GJ. Conventional laboratory housing increases morbidity and mortality in research rodents: results of a meta-analysis. BMC Biol. 2022;20: 15. doi: 10.1186/s12915-021-01184-0.

7. Prager EM, Bergstrom HC, Grunberg NE, Johnson LR. The importance of reporting housing and husbandry in rat research. Front Behav Neurosci. 2011;5:38. doi: 10.3389/fnbeh.2011.00038.

8. Bradshaw AL, Poling A. Choice by rats for enriched versus standard home cages: plastic pipes, wood platforms, wood chips and paper towels as enrichment items. J Exp Anal Behav. 1991;55(2):245–250. doi: 10.1901/jeab.1991.55-245.

9. Mellor DJ, Beausoleil NJ, Littlewood KE, McLean AN, McGreevy PD, Jones B, Wilkins C. The 2020 five domains model: Including human-animal interactions in assessments of animal welfare. Animals. 2020;10(10):1870. doi: 10.3390/ani10101870.

10. Miller LJ, Vicino GA, Sheftel J, Lauderdale LK. Behavioral diversity as a potential indicator of positive welfare. Animals. 2020;10(7):1211. doi:10.3390/ani10071211.

11. Canadian Council on Animal Care (CCAC) CCAC guidelines: Rats. 2020. Available from: https://ccac.ca/Documents/Standards/Guidelines/CCAC_Guidelines_Rats.pdf

12. Gärtner K, Büttner D, Döhler K, Friedel R, Lindena J, Trautschold I. Stress response of rats to handling and experimental procedures. Lab Anim. 1980;14:267–274. doi:10.1258/002367780780937454.

13. Costa R, Tamascia ML, Nogueira MD, Casarini DE, Marcondes FK. Handling of adolescent rats improves learning and memory and decreases anxiety. J Am Assoc Lab Anim Sci. 2012; 51(5): 548–553.

14. Schmitt U, Hiemke C. Strain differences in open-field and elevated plus-maze behavior of rats without and with pretest handling. Pharmacol Biochem Behav. 1998;59(4):807–811. doi:10.1016/S0091-3057(97)00502-9.

15. Manseur A, Bairi A, Bakeche A, Djouini A, Tahraoui A. Effect of handling by human being neonatal period on anxiety and depression-like behavior of adult rats. Adv Anim Vet Sci. 2019;7(12):1113–1119.

16. Hirsjärvi PA, Junnila MA, Väliaho TU. Gentled and non-handled rats in a stressful open-field situation; differences in performance. Scand J Psychol. 1990;31:259–265. doi:10.1111/j.1467-9450.1990.tb00838.x.

17. Maurer BM, Döring D, Scheipl F, Küchenhoff H, Erhard MH. Effects of a gentling programme on the behaviour of laboratory rats towards humans. Appl Anim Behav Sci. 2008;114:554–571. doi:10.1016/j.applanim.2008.04.013.

18. Costa R, Tamascia ML, Nogueira MD, Casarini DE, Marcondes FK. Handling of adolescent rats improves learning and memory and decreases anxiety. J Am Assoc Lab Anim Sci. 2012; 51(5): 548–553.

19. Davis H, Pérusse R. Human-based social interaction can reward a rat’s behavior. Anim Learn Behav.1988;16(1): 89–92. doi:10.3758/BF03209048.

20. Gärtner K, Büttner D, Döhler K, Friedel R, Lindena J, Trautschold I. Stress response of rats to handling and experimental procedures. Lab Anim. 1980;14:267–274. doi:10.1258/002367780780937454.

21. Sorge RE, Martin LJ, Isbester KA, Sotocinal SG, Rosen S, Tuttle AH, Wieskopf JS, Acland EL, Dokova A, Kadoura B, Leger P, Mapplebeck JCS, McPhail M, Delaney A, Wigerblad G, Schumann AP, Quinn T, Frasnelli J, Svensson CI, Sternber WF, Mogil JS. Olfactory exposure to males, including men, causes stress and related analgesia in rodents. Nat Methods. 2014;11(6):629–632. doi:10.1038/nmeth.2935.

22. Herzog HA. Gender differences in human-animal interactions: a review. Anthrozoos. 2007;20:7–21. doi:10.2752/089279307780216687.

23. Pellow S, Chopin P, File SE, Briley M. Validation of open:close arm entries in the elevated plus-maze as a measure of anxiety in the rat. J Neurosci Methods. 1985;14:149–167. doi: 10.1016/0165-0270(85)90031-7.

24. Feyissa DD, Aher YD, Engidawork E, Höger H, Lubec G, Korz V. Individual differences in male rats in a behavioral test battery: A multivariate statistical approach. Front Behav Neurosci. 2017;11:26. doi:10.3389/fnbeh.2017.00026.

25. Handley SL, Mithani S. Effects of alpha-adrenoceptor agonists and antagonists in a maze-exploration model of ‘fear’-motivated behaviour. Naunyn Schmiedebergs Arch Pharmacol. 1984;327:1–5. doi:10.1007/BF00504983.

26. Walf AA, Frye CA. The use of elevated plus maze as an assay of anxiety-related behavior in rodents. Nat Protoc. 2007;2:322–328. doi: 10.1038/nprot.2007.44.

27. Abou-Ismail UA, Burman OHP, Nicol CJ, Mendl M. The effects of enhancing cage complexity on the behaviour and welfare of laboratory rats. Behavi Process. 2010;85(2): 172–180. doi: 10.1016/j.beproc.2010.07.002.

28. Pinelli C, Leri F, Turner PV. 2017. Long term physiologic and behavioural effects of housing density and environmental resource provision for adult male and female Sprague Dawley rats. Animals. 7(6):E44. doi: 10.3390/ani7060044. http://www.mdpi.com/2076-2615/7/6/44/pdf

29. Modlinska K, Chrzanowska A, Pisula W. The impact of changeability of enriched environment on exploration in rats. Behav Processes. 2019;164:78–85. doi:10.1016/j.beproc.2019.04.015.

30. Blanchard RJ, McKittrick CR, Blanchard DC. Animal models of social stress: effects on behavior and brain neurochemical systems. Physiol Behav. 2001;73(3): 261–271. doi: 10.1016/S0031-9384(01)00449-8.

31. Pinzón-Parra C, Vidal-Jiménez B, Camacho-Abrego I, Flores-Gómez AA, Rodríguez-Moreno A, Flores G. Juvenile stress causes reduced locomotor behavior and dendritic spine density in the prefrontal cortex and basolateral amygdala in Sprague-Dawley rats. Synapse. 2018;73:e22066. doi:10.1002/syn.22066.

32. Mellor DJ. Enhancing animal welfare by creating opportunities for positive affective engagement. N Z Vet J. 2014;63(1):3–8. doi:10.1080/00480169.2014.926799.

33. Jirkof P, Rudeck J, Lewejohann L. Assessing affective state in laboratory rodents to promote animal welfare—What is the progress in applied refinement research? Animals. 2019;9(12):1026. doi:10.3390/ani9121026.

34. Kalueff AV, Tuohimaa P. The grooming analysis algorithm discriminates between different levels of anxiety in rats: potential utility for neurobehavioral stress research. J Neurosci Methods. 2005;143(2):169–177. doi:10.1016/j.jneumeth.2004.10.001.

35. Estanislau C, Veloso AWN, Filgueiras GB, Maio TP, Dal-Cól MLC, Cunha DC, Klein R, Carmona LF, Fernández-Teruel A. Rat self-grooming and its relationships with anxiety, derousal and perseveration: evidence for a self-grooming trait. Physiol Behav. 2019;209:112585. doi:10.1016/j.physbeh.2019.112585.

36. Büttner D. Upright standing in the laboratory rat – time expenditure and its relation to locomotor activity. J Exp Anim Sci. 1993;36(1): 19–26.

37. Vachon P. Double decker enrichment cages have no effect on long term nociception in neuropathic rats but increase exploration while decreasing anxiety-like behaviors. Scand J Lab Anim Sci. 2014;40:1–6.

38. Wheeler RR, Swan MP, Hickman DL. Effect of multilevel laboratory rat caging system on the well-being of the singly-housed Sprague-Dawley rat. Lab Anim. 2015;49:10–19. doi:10.1177/0023677214547404.

39. Dodelet-Devillers A, Zullian C, Beaudry F, Gourdon J, Chevrette J, Hélie P, Vachon P. Physiological and pharmacokinetic effects of multilevel caging on Sprague Dawley rats under ketamine-xylazine anesthesia. Exp Anim. 2016;65(4):383–392. doi:10.1538/expanim.16-0026.

40. Mieske P, Hobbiesiefken U, Fischer-Tenhagen C, Heinl C, Hohlbaum K, Kahnau P, Meier J, Wilzopolski J, Butzke D, Rudeck J, Lewejohann L, Diederich K. Bored at home? A systematic review on the effect of environmental enrichment on the welfare of laboratory rats and mice. Front Vet Sci. 2022;9:899219. doi:10.3389/fvets.2022.899219.

41. Browning H, Weit W. Freedom and animal welfare. Animals. 2021;11: 1148. doi: 10.3390/ani11041148.

42. Bayne K, Turner PV. Animal welfare standards and international collaborations. ILAR J. 2019;60(1):86–94. doi:10.1093/ilar/ily024.

43. Boguszewski P, Zagrodzka J. Emotional changes related to age in rats—a behavioral analysis. Behav Brain Res. 2002;133(2):323–332. doi: 10.1016/s0166-4328(02)00018-9.

44. Kondrakiewicz K, Kostecki M, Szadzińska W, Knapska E. Ecological validity of social interaction tests in rats and mice. Genes Brain Behav. 2019;18(1):e12525. doi:10.1111/gbb.12525.

45. Martin B, Ji S, Maudsley S, Mattson MP. "Control" laboratory rodents are metabolically morbid: why it matters. Proc Natl Acad Sci U S A. 2010;107(14):6127–6133. doi:10.1073/pnas.0912955107.

46. Wei Y, et al. Enriched environment-induced maternal weight loss reprograms metabolic gene expressions in mouse offspring. J Biol Chem. 2015;290(8):4604–4619. doi:10.1074/jbc.M114.605642.

47. Smith BL, et al. Behavioral and physiological consequences of enrichment loss in rats. Psychoneuroendocrinology. 2017;77:37–46. doi:10.1016/j.psyneuen.2016.11.040.

48. Morano R, Hoskins O, Smith BL, Herman JP. Loss of environmental enrichment elicits behavioral and physiological dysregulation in female rats. Front Behav Neurosci. 2019;12:287. doi:10.3389/fnbeh.2018.00287.

49. Mazhary H, Hawkins P. Applying the 3Rs: a case study on evidence and perceptions relating to rat cage height in the UK. Animals. 2019;9:1102. doi:10.3390/ani9121104.

50. Rault JL, Waiblinger S, Boivin X, Hemsworth P. The power of a positive human-animal relationship for animal welfare. Front Vet Sci. 2020;7:590867. doi: 10.3389/fvets.2020.590867.

51. Reynolds S, Lane SJ, Richards L. Using animal models of enriched environments to inform research on sensory integration intervention for the rehabilitation of neurodevelopmental disorders. J Neurodevelop Disord. 2010;2:120–132. doi:10.1007/s11689-010-9053-4.

52. Baratta MV, Seligman MEP, Maier SF. From helplessness to controllability: toward a neuroscience of resilience. Front Psychiatry. 2023;14:1170417. doi:10.3389/fpsyt.2023.1170417.

53. Silveira KM, Joca S. Learned Helplessness in Rodents. In: Neuromethods. Springer US; 2022. p. 161–184. doi:10.1007/978-1-0716-2748-8_9.

54. Brydges NM, Leach M, Nicol K, Wright R, Bateson M. Environmental enrichment induces optimistic cognitive bias in rats. Anim Behav. 2011;81(1):169–175. doi:10.1016/j.anbehav.2010.09.030.

55. Burman OH, Parker RM, Paul ES, Mendl MT. Anxiety-induced cognitive bias in non-human animals. Physiol Behav. 2009;98(3):345–350. doi:10.1016/j.physbeh.2009.06.012.

56. Tsang B, Berlai R. Researchers, animal support and regulatory staff: symbiosis or antagonism? Lab Anim Res. 2022;38:19. doi: 10.1186/s42826-022-00129-0.

57. O’Malley CI, Hubley R, Moody C, Turner PV. Use of nonaversive handling and training procedures for laboratory mice and rats: Attitudes of American and Canadian laboratory animal professionals. Front Vet Sci. 2022;9:1040572. doi:10.3389/fvets.2022.1040572.

58. Freedman LP, Cockburn IM, Simcoe TS. The economics of reproducibility in preclinical research. PLoS Biol. 2018;16(4):e1002626. doi:10.1371/journal.pbio.1002626.

59. Percie du Sert N, et al. PLoS Biology. 2020 Jul;18(7):e3000410. doi: 10.1371/journal.pbio.3000410.

60. Baumans V, Van Loo PLP, Pham TM. Standardisation of environmental enrichment for laboratory mice and rats: utilization, practicality and variation in experimental results. Scand J Lab Anim Sci. 2010;37(2): 101–114. doi: 10.23675/sjlas.v37i2.209.

61. LaFollette MR, Riley MC, Cloutier S, Brady CM, O’Haire ME, Gaskill BN. Laboratory animal welfare meets human welfare: A cross-sectional study of professional quality of life, including compassion fatigue in laboratory animal personnel. Front Vet Sci. 2020;7:114. doi:10.3389/fvets.2020.00114.

62. Murray J, Bauer C, Vilminot N, Turner PV. Strengthening workplace well-being in research animal facilities. Front Vet Sci. 2020;7:573106. doi:10.3389/fvets.2020.573106.

